# Dietary Amino Acids Impact LRRK2-induced Neurodegeneration in Parkinson’s Disease Models

**DOI:** 10.1101/2020.01.13.905471

**Authors:** Vinita G. Chittoor-Vinod, Steffany Villalobos-Cantor, Hanna Roshak, Kelsey Shea, Ian Martin

**Author notes:** Corresponding author:* Ian Martin, PhD, Department of Neurology, Oregon Health and Science University. 3181 SW Sam Jackson Park Road, Portland, OR 97239.

## Abstract

The G2019S mutation in leucine-rich repeat kinase 2 (LRRK2) is a common cause of Parkinson’s disease (PD) and results in age-related dopamine neuron loss and locomotor dysfunction in *Drosophila melanogaster* through an aberrant increase in bulk neuronal protein synthesis. Under non-pathologic conditions, protein synthesis is tightly controlled by metabolic regulation. Whether nutritional and metabolic influences on protein synthesis can modulate the pathogenic effect of LRRK2 on protein synthesis and thereby impact neuronal loss is a key unresolved question. Here, we show that LRRK2 G2019S-induced neurodegeneration is critically dependent on dietary amino acid content. Low dietary amino acid concentration prevents aberrant protein synthesis and blocks LRRK2 G2019S-mediated neurodegeneration in *Drosophila* and rat primary neurons. Unexpectedly, a moderately high amino acid diet also blocks dopamine neuron loss and motor deficits in *Drosophila* through a separate mechanism involving stress-responsive activation of 5’-AMP-activated protein kinase (AMPK) and neuroprotective induction of autophagy, implicating the importance of protein homeostasis to neuronal viability. At the highest amino acid diet of the range tested, PD-related neurodegeneration occurs in an age-related manner, but is also observed in control strains, suggesting that it is independent of mutant LRRK2 expression. We propose that dietary influences on protein synthesis and autophagy are critical determinants of LRRK2 neurodegeneration, opening up possibilities for future therapeutic intervention.

## INTRODUCTION

Parkinson’s disease (PD) pathology is clearly age-related, leading to the hypothesis that PD and aging manifest through common mechanisms (Collier *et al*. 2017). Dietary restriction, i.e. a moderate reduction in food intake, impacts aging in many species including primates (Gems and Partridge 2013). Evidence in animal models and more recently humans indicates that the beneficial effects of dietary restriction are primarily attributable to reduced protein intake (Kapahi *et al*. 2004; Grandison *et al*. 2009; Levine *et al*. 2014). Dietary protein influences protein synthesis through governing the pool of available amino acids and by signaling to metabolic pathways that couple nutrient sensing to anabolic output. A central amino acid sensor in eukaryotes is the mechanistic target of rapamycin complex 1 (mTORC1) which regulates protein synthesis in response to the supply of specific amino acids (Kim *et al*. 2002). mTORC1 is evolutionarily conserved in invertebrates such as *Drosophila,* where the orthologous complex is called TORC1 (Oldham 2011). Blocking TORC1 signaling in flies by manipulating the pathway components dTsc1, dTsc2, dTOR or S6K phenocopies the life span-extending effects of dietary restriction (Kapahi *et al*. 2004). Another important effector of TORC1 is the protein synthesis-repressor 4E-BP (eIF4E-binding protein), whose activity is potentiated upon TORC1 inhibition through dietary restriction. Silencing 4E-BP prevents the maximal life span extension obtained through dietary restriction, providing direct evidence that enhanced longevity is dependent on TORC1 regulation of protein synthesis (Zid *et al*. 2009). Despite a link with aging, the role of protein consumption and protein synthesis in age-related neurodegeneration is largely unexplored. Studies in mice and primate toxin models of PD show that restricted caloric intake leads to attenuated motor deficits and nigral dopamine neuron loss (Maswood *et al*. 2004; Bayliss *et al*. 2016), but no studies to date have investigated the potential for altered protein intake to impact PD-related phenotypes.

A broad role for leucine-rich repeat kinase 2 (LRRK2) mutations in familial and idiopathic PD has emerged (Healy *et al*. 2008; Simon-Sanchez *et al*. 2009). Expression of the common pathogenic mutation LRRK2 G2019S causes robust age-related and selective loss of dopamine neurons, with accompanying locomotor deficits in *Drosophila* (Liu *et al*. 2008). These phenotypes contrast with existing rodent transgenic or knock-in LRRK2 models which typically display more subtle PD-related phenotypes without overt neuron loss (Herzig *et al*. 2011), highlighting the utility of *Drosophila* models as an important tool for understanding LRRK2-induced neurodegeneration. We and others previously showed that expressing LRRK2 G2019S in flies causes elevated neuronal bulk protein synthesis, and that blocking this through protein synthesis inhibitors ameliorates age-related dopamine neuron loss and locomotor dysfunction (Imai *et al*. 2008; Martin *et al*. 2014b; Penney *et al*. 2016). The observation that elevated bulk protein synthesis contributes to PD-related phenotypes in LRRK2 G2019S flies generates a compelling need to understand if metabolic regulation of protein synthesis through diet can impact dopamine neuron survival with age. Recently, a chemically-defined “holidic” *Drosophila* food medium was described that contains all dietary components necessary to promote optimal fly survival and fecundity (Piper *et al*. 2014). Holidic food is a completely defined and synthetic medium composed of purified ingredients which contrasts with oligidic medium which is yeast-based and contains natural ingredients but is not amenable to modifying individual nutrient classes.

Here, we used a holidic *Drosophila* diet approach to investigate the impact of dietary amino acid levels on PD-related phenotypes in LRRK2 G2019S *Drosophila*. Upon testing a range of amino acid concentrations, we found that substantially reducing amino acid concentration relative to the estimated content of our standard chemically-undefined *Drosophila* food rescues age-related dopamine neuron loss and locomotor dysfunction in LRRK2 G2019S flies. This is associated with a reduction in bulk protein synthesis, but independent of TORC1 activity, autophagy induction or activation of the sensor of amino acid deficiency GCN2, suggesting that this level of restriction does not constitute amino acid starvation. Elevating dietary amino acid levels within a range unexpectedly suppresses PD-related phenotypes through a distinct mechanism involving stress-responsive induction of AMPK with associated enhancement of autophagy and dampening of TORC1 activity in aged flies. We further extend our findings to show that restriction of amino acids rescues neurite shortening and cell death associated with LRRK2 G2019S expression in rat cortical neurons. Collectively, our results uncover an exquisite sensitivity of LRRK2 G2019S-mediated neurodegeneration to dietary amino acid content in *Drosophila* and rodent neurons.

## MATERIALS AND METHODS

### DROSOPHILA STOCKS AND CULTURE

The *UAS-LRRK2-G2019S* and *UAS-LRRK2-WT* lines were a gift from Wanli Smith and were backcrossed to *w^1118^* as described elsewhere (Martin *et al*. 2014b). Expression of these transgenes was driven by the dopaminergic *Ddc-GAL4* driver (BDSC 7010) or the pan-neuronal *elav-GAL4* driver (BDSC 8765) as indicated. *GAL4* drivers were additionally crossed to *w^1118^* to generate non-LRRK2 transgenic controls. RNAi fly stocks used were *UAS-AMPK-IR 1* (BDSC 25931); *UAS-AMPK-IR 2* (BDSC 32371) and *UAS-atg1-IR* (BDSC 35177). *UAS-atg1^DN^* (*UAS-atg1^KQ#5B^*) is a dominant negative form of *atg1* (Scott *et al*. 2007) and was a gift from Eric Baehrecke. For LRRK2 G2019S epistasis experiments, double-transgenic *Ddc-GAL4; UAS-LRRK2-G2019S* and *Ddc-GAL4; UAS-LRRK2-WT* lines were generated. All flies were maintained at 25°C and 60% relative humidity under a 12h:12h light:dark cycle.

### HOLIDIC MEDIA PREPARATION

Chemically-defined holidic media were prepared following detailed protocols described by Piper *et al*. (Piper *et al*. 2014) and using the HUNTaa composition of amino acids. See Table S1 for specific formulations of each holidic medium used. All food components were from the same sources as used in Piper et al. and following preparation, holidic media were stored at 4 °C until use. The total amino acid-based nitrogen content (millimolar, designated as “N”) of the four diets used were: 25N, 50N, 200N and 400N. This was achieved by modifying the levels of all essential and non-essential amino acids equally while keeping all other dietary components constant. Sucrose levels were standardized to 50S in all media as per Piper et al. in order to promote optimal fly longevity. Flies were reared on our standard chemically-undefined medium and transferred to holidic media as adults to avoid potential effects of holidic medium or altered amino acid levels on development, as previously reported (Piper *et al*. 2014). Cohorts (0-3 days-old, selecting against flies with visible signs of recent eclosion) were collected under brief anesthesia and transferred to 25N, 50N, 200N or 400N holidic food diets at 25 flies/vial. Flies were housed on a single holidic diet and switched to fresh food every 3 days until testing. For the 25N-F/H/W medium used to assess protein synthesis, phenylalanine (F), histidine (H) and tryptophan (W) were present at 25N medium levels, while all other amino acids were at 50N levels (see Table S3 for composition).

### ESTIMATION OF PROTEIN NITROGEN MOLARITY IN STANDARD DROSOPHILA MEDIUM

The standard chemically-undefined medium used in prior studies (Martin *et al*. 2014b) contains (per liter) yeast 15 g, cornmeal 60 g, agar 5 g, corn syrup 67 ml, propionic acid 4 ml, tegosept 2.4 ml. Principle sources of protein in this medium are brewer’s yeast (15 g/L; ∼40% protein content) and yellow cornmeal (60 g/L; ∼7% protein content). An estimate of protein-derived nitrogen molar content (N) was derived from total protein (10.2 g/L) via multiplying by 0.16 (Jones factor) and then dividing by the molar mass of nitrogen (14 g/mol) = 117 mM N.

### FOOD INTAKE

Blue dye (erioglaucine disodium salt; Sigma-Aldrich, 861146) was added at 1.5% (w/v) to holidic media during food preparation to assess food intake based on established procedures (Bjordal *et al*. 2014). Female flies (n=10 per condition) were transferred to blue dye-labeled holidic medium at the same amino acid level as the medium they had been housed on and kept at 25°C (60% relative humidity) for 2 hours. The experiment was terminated by transferring the flies into microtubes and flash freezing in liquid nitrogen. Whole bodies were washed and lysed in ice-cold water. Supernatant was collected after centrifugation at 13,000 rpm for 5 mins at 4°C. The amount of ingested blue dye in the supernatant was measured spectrophotometrically at 620 nm (Fisher Scientific Multiskan FC). Sample readings were normalized to the number of flies.

### FLY BRAIN IMMUNOHISTOCHEMISTRY AND DOPAMINE NEURON COUNTS

Brains (12-15 per condition) from flies aged on holidic media for 6 weeks were harvested, fixed and permeabilized (4% PFA in PBS-T pH 7.4). Fixed brains were blocked in 5% normal donkey serum (Sigma-Aldrich, D9663) for 1 h at room temperature (RT), followed by incubation with anti-tyrosine hydroxylase (1:1000; Immunostar, 22941) for 72 hours at 4°C followed by extensive washing and incubation with Alexa-fluor 488 secondary antibody (1:2000; Thermo Fisher Scientific, A-21202) for 72 hours at 4°C. Brains were washed extensively and whole-mounted in SlowFade Gold Antifade mounting medium (Thermo Fisher Scientific, S36938). Confocal z-stack images of the stained brains were acquired on Zeiss LSM 780 at 1 µm slice intervals. Projection images of the protocerebral posterior lateral 1 (PPL1) cluster were used for dopamine neuron counts.

### NEGATIVE GEOTAXIS ASSAY

Cohorts of 100 female flies (25 flies/vial) aged on holidic media were tested for motor function using negative geotaxis assays performed and analyzed as described elsewhere (Gargano *et al*. 2005). The performance of flies in a single vial was calculated from the average height climbed by all flies in the vial 4-seconds after initiating climbing behavior. This generated a single datum (N =1) and the average scores of 4 vials were tested per line (N = 4).

### ^35^S-METHIONINE/CYSTEINE LABELING OF DROSOPHILA BRAINS

*Ex vivo* labeling of protein synthesis in fly brains was carried out based on previously reported methods (Essers *et al*. 2016). Fly brains (8 per genotype) were harvested in Schneider’s medium on ice and transferred to Schneider’s medium containing ^35^S-methionine/cysteine (2 µCi/µl; Perkin Elmer Labs Inc., NEG709A) and 30 µM chloroquine (to inhibit autophagy). Brains were incubated for 20 min at 25 °C and washed twice in chilled PBS. On the final wash, all PBS was removed and the brains were flash frozen. Brains were homogenized by pestle and mortar in RIPA extraction buffer on ice. Protein from 50% of the lysate was precipitated by the addition of methanol and heparin (lysate:heparin(100mg/ml):methanol volume ratio of 150:1.5:600), centrifuged at 14,000x g for 2 minutes, and supernatant was removed and the pellet air dried. The protein pellet was resuspended in 8M urea/150 mM Tris, pH 8.5 and ^35^S-methionine/cysteine incorporation was measured by scintillation counting and normalized to the number of brains. A portion of the remaining RIPA extraction lysate was separated by SDS-PAGE with equal protein loading, and transferred to nitrocellulose membrane. The membrane was dried and ^35^S-methionine/cysteine incorporation was visualized by exposing to a phosphorimaging cassette followed by scanning using a Typhoon scanner.

### WESTERN BLOTTING

Fly heads (25 flies per condition) were harvested at the indicated ages and extracted in a 1% NP-40 lysis buffer: 50 mM Tris-HCl pH 7.5, 150 mM NaCl, 5 mM EGTA, 1% (v/v) NP40, 100 mM NaF, 2 mM β-glycerophosphate, 2% (v/v) phosphatase inhibitor (Sigma-Aldrich, P0044) and protease inhibitors. Sample protein concentrations were determined by BCA assay (Thermo Fisher Scientific, PI3225) and equal total protein amounts were electrophoretically separated by SDS-PAGE and transferred to nitrocellulose membranes (Thermo Fisher Scientific, PI88018). Blots were incubated with primary antibody for 18 hours at 4°C. Primary antibodies used for Western blotting include: p-S6K (1:750; Cell Signaling Technology, 9209); p-4EBP (1:750; Cell Signaling Technology, 2855); Non-p-4EBP (1:1000; Cell Signaling Technology, 4923); Actin (1:1000, Cell Signaling Technology, 4967); p-LRRK2 (1:750; Abcam, ab203181); LRRK2 (1:1000; Cell Signaling Technology, 13046); Rps15 (1:500; Sigma-Aldrich, WH0006209M1); p-AMPK (1:1000; Cell Signaling Technology, 2535); AMPK (1:500; Millipore, MABS1232) and LAMP1 (1:1000; Abcam, ab30687). A P-RPS15 antibody used was previously described ^23^. Blots were imaged using a Syngene G:BOX. Semi-quantitative densitometric analyses were performed using ImageJ software (NIH).

### QUANTITATIVE RT-PCR

Total RNA was isolated from fly heads (n=15 per condition) using TRIzol reagent (Thermo Fisher Scientific, 15596026) and quantified on NanoDrop 2000c (Thermo Scientific). 100 ng total RNA was reverse transcribed using Superscript IV first-strand synthesis system (Thermo Fisher Scientific, 18091050), using random hexamers. Quantitative real-time PCR assays were performed in Maxima SYBR Green/ROX qPCR master mix (Thermo Fisher Scientific, K0222) using a Thermo Fisher Scientific QuantStudio 3 Real Time PCR system under the following sequence of conditions: 50°C for 2 mins, 95°C for 10 mins then 40 cycles (c) of 15 s at 95°C and 1 min at variable hybridization temperature (H). For grim transcript detection, cycle parameters described elsewhere (Zheng *et al*. 2010) were followed. Primer sequences used in this study are as follows: dSesn: (F) 5’-CTCGACTCGATCCCTCCG-3’; (R) 5’-CAGGTCATCGAGCTCGTCC-3’; c=40; H= 58°C; atg8a: (F) 5’-ACCAGGAACATCACGAGGAG-3’; (R) 5’-TTAGCTAACTCGCCGTCCAT-3’; c= 40; H= 58°C; atg9: (F) 5’-TCGTCTGGCTACTTGCCTTT-3’; (R) 5’-AACCAGGTACGTGACCCAAG-3’; c= 40; H= 58°C; grim: (F) 5’-GTCGTCCTCATCGTTGTTCTGAC-3’; (R) 5’-CCATCGCCTATTTCATACCCG-3’; c= 40; H= 57.9°C; reaper: (F) 5’-AGTCACAGTGGAGATTCCTGG-3’; (R) 5’-TGCGATATTTGCCGGACTTTC-3’; c=40; H=57°C; actin: (F) 5’-GTGAAATCGTCCGTGACATC-3’; (R) 5’-GGCAGCTCGTAGGACTTCTC-3’; c= 58°C; H=40.

### RAT CORTICAL NEURON CULTURE

E18 Sprague Dawley rat cortices (BrainBits LLC) were dissociated in trypsin substitute TrypLE enzyme mix (Fisher, 12-605-010) for 6 min at 37 °C and triturated vigorously with a silanized glass pipet for 1 min. After allowing debris to settle, the supernatant containing dissociated cells was centrifuged at 120 x g for 1 min, resuspended in Neurobasal without phenol red (Fisher, 12-348-017)/1% B27 supplement (Fisher, 17-504-044)/ 1% glutaMAX (Fisher, 35-050-061) and plated at a density of 5.0×10^4^ cells per well on poly-D-Lysine (Sigma, P6407) coated 24 well glass-bottom plates (Fisher, 50-114-9070). Neurons were cultured at 37°C/5% CO_2_ and glial growth was inhibited at DIV4 by addition of 5-fluoro-2’-deoxyuridine (30 μM)(Sigma, F0503). A 50% fresh culture medium change was performed every 4 days.

### GENERATION OF AMINO ACID-DILUTED RAT NEURON CULTURE MEDIUM

A neurobasal-based medium without amino acids was generated using the formulation of standard neurobasal without phenol red (Fisher, 12-348-017) and omitting all amino acids (Table S2). All other standard neurobasal components were present at standard formula concentrations and using cell culture-grade reagents. This medium was vacuum filtered and combined with standard neurobasal medium in order to dilute amino acids to a range of 0.4-0.8 x relative to standard neurobasal medium. All media were then combined with B27 supplement (but no GlutaMAX for L-glutamine as this is an additional source of amino acid) and filtered again.

### LRRK2 RAT NEURON TOXICITY ASSAYS

Toxicity (loss of neurites and appearance of TUNEL-positive nuclei) in rat cortical neurons overexpressing LRRK2 WT or LRRK2 G2019S was assessed using previously described methods (Martin *et al*. 2014b) 48 h following plasmid transfection using Lipofectamine 2000 (Fisher, 11-668-027). At DIV 10, neurons were switched to Opti-MEM (Gibco, 115439) reduced serum medium and transfected with plasmid DNA. Myc tagged LRRK2 and pEGFP(N3) for tracing neurites at a 10:1 ratio were mixed with Lipofectamine 2000 at a 1:3 ratio in Opti-MEM. Transfection complex was removed after 2 hours and replaced with amino acid-modified or standard culture medium (0.4-1.0 dilution, see above for description of amino acid diluted media) for an additional 48 h of culture. Neurons were washed, fixed (4% PFA in PBS-T pH 7.4), blocked and permeabilized for 1 h in 10% donkey serum/PBS-T/0.3% triton-X-100 pH 7.4 and incubated overnight (4 °C) in mouse anti-GFP (3E6) (Thermo A-11120) and rabbit anti-LRRK2 (D18E12) (CST, 13046) primary antibodies, washed extensively, and incubated in anti-mouse Alexa-Fluor 488 (1:2000; Fisher, A21202) and anti-rabbit Alexa-Fluor 647 (1:2000, Fisher, A32733) secondary antibodies for 1 h at RT. Cell death was assessed by TUNEL assay (with TMR, red-dUTP, Sigma, 12156792910) and nuclei were labeled with DAPI stain (Molecular Probes, D1306) following manufacturer’s protocols. Neurons were imaged on a Zeiss LSM 780 confocal microscope. The vast majority of GFP-positive neurons also overexpressed LRRK2 allowing the effects of LRRK2 expression to be visualized in these neurons. Injured neurons without at least one smooth extension (neurite) twice the length of the cell body were recorded and the percentage of GFP-positive injured neurons in each experimental group relative to all GFP-positive neurons was derived. The percentage of GFP-positive neurons with TUNEL-positive nuclei was also measured. At least 75 neurons were counted per group per independent experiment.

### PROTEIN SYNTHESIS SUnSET ASSAY IN RAT NEURONS

The surface sensing of translation (SUnSET) assay permits the non-radioactive monitoring of global protein synthesis in mammalian cells by visualization of low concentration puromycin incorporation into nascent polypeptide chains (Schmidt *et al*. 2009). Puromycin incorporation results in nascent chain termination but when administered at low micromolar concentrations does not substantially interfere with global protein synthesis. DIV10 neurons were cultured in amino acid-modified (see above for description) or standard culture medium for 48 h post-transfection and then exposed to 2 μM puromycin (Millipore, 540222) for 10 min at 37°C/5% CO_2_. Following incubation, neurons were immediately fixed (4% PFA in PBS-T pH 7.4), blocked and permeabilized for 1 h in 10% donkey serum/PBS-T/0.3% triton-X-100 pH 7.4 and incubated for 2 h (RT) in mouse anti-puromycin 12D10 antibody (Sigma Aldrich, MABE343), washed extensively, then incubated in anti-mouse Alexa-fluor 488 secondary antibody (1:2000, Fisher, A21202) for 1 h at RT. Neurons were imaged on a Zeiss LSM 780 confocal microscope. Immunofluorescence signal (integrated density counts of cells corrected for background) was quantified using Image J where a single n represents total signal from 20 neurons.

### ADP/ATP RATIO ASSAY

The ADP/ATP ratio assay kit (Sigma MAK135) was used following all manufacturer instructions. Fly heads were homogenized in ATP reagent and centrifuged for 10 min at 4 °C. Luminescence in the supernatant was read using a SpectraMax microplate reader and then the ADP reagent was added to assess ADP levels and derive ADP/ATP ratio according to manufacturer instructions.

### EXPERIMENTAL DESIGN AND STATISTICAL ANALYSES

Quantified data presented in all figures are mean ± SEM. Statistical analysis details for each individual experiment are described in Figure legends, including number of flies or number of groups of flies used (N), statistical tests (one-way or two-way ANOVA), and Bonferroni post hoc analysis with associated p values. All statistical analyses were performed using GraphPad Prism software.

## RESULTS

### Dietary amino acids regulate PD-related phenotypes in *Drosophila*

Flies expressing pathogenic human LRRK2 G2019S via the dopaminergic *Ddc-GAL4* driver display an age-related loss of dopamine neurons with accompanying locomotor dysfunction at 6 weeks of age (Liu *et al*. 2008; Martin *et al*. 2014b). Flies expressing human wild-type LRRK2 (LRRK2 WT) do not exhibit these phenotypes, indicating that they arise from mutation-specific effects. Considering the established contribution of elevated bulk protein synthesis to LRRK2 G2019S-induced neurodegeneration (Imai *et al*. 2008; Martin *et al*. 2014b; Penney *et al*. 2016), we sought to determine whether dietary influences on protein synthesis can impact neurodegeneration. We aged flies expressing human LRRK2 WT or LRRK2 G2019S on a range of holidic diets in which the concentration of all amino acids is varied while all other food constituents remain constant throughout adulthood (Table S1). In these diets, amino acid content is denoted by the total amino acid-derived millimolar nitrogen concentration (N) of the food (Piper *et al*. 2014). Amino acid and other nutrient content in the holidic medium is fixed and defined which contrasts to those of a “standard” Drosophila laboratory diet, where it has shown to vary widely between labs (Piper *et al*. 2014) and possibly also between batches of food within the same lab, based on lot-to-lot ingredient variability. Piper et al., previously demonstrated that aging adult flies on diets with an amino acid concentration range between 0N and 400N causes major effects on median life span, which peaks on 50N-100N amino acid diets and dramatically shortens at both extremes of this amino acid range. To assess if a similar range of dietary amino acids can impact PD-related phenotypes caused by mutant LRRK2, we generated diets varying in amino acid concentrations from 25N to 400N and examined dopamine neuron loss and motor deficits following 6 weeks of aging on these diets. Both dopamine neuron viability and locomotor function are substantially affected by dietary amino acid concentration at this age (Fig. 1). Strikingly, LRRK2 G2019S-dependent dopamine neuron death and locomotor dysfunction are only observed in flies on a 50N amino acid diet. LRRK2 G2019S flies aged on 25N or 200N amino acid diets exhibit a rescue of dopamine neuron viability and motor abilities relative to those on a 50N diet (Fig. 1). The 400N diet elicits a loss of dopamine neurons which occurs in all fly strains tested, suggesting that it is LRRK2-independent (Fig. 1a,b). There is no significant effect of dietary amino acid levels on locomotor function in young control or mutant LRRK2 flies housed on these diets for one week (Fig. S1a), suggesting that the diet effects on neurodegenerative phenotypes seen in aged LRRK2 G2019S flies on a 50N diet are age-dependent.

**Figure. 1.**
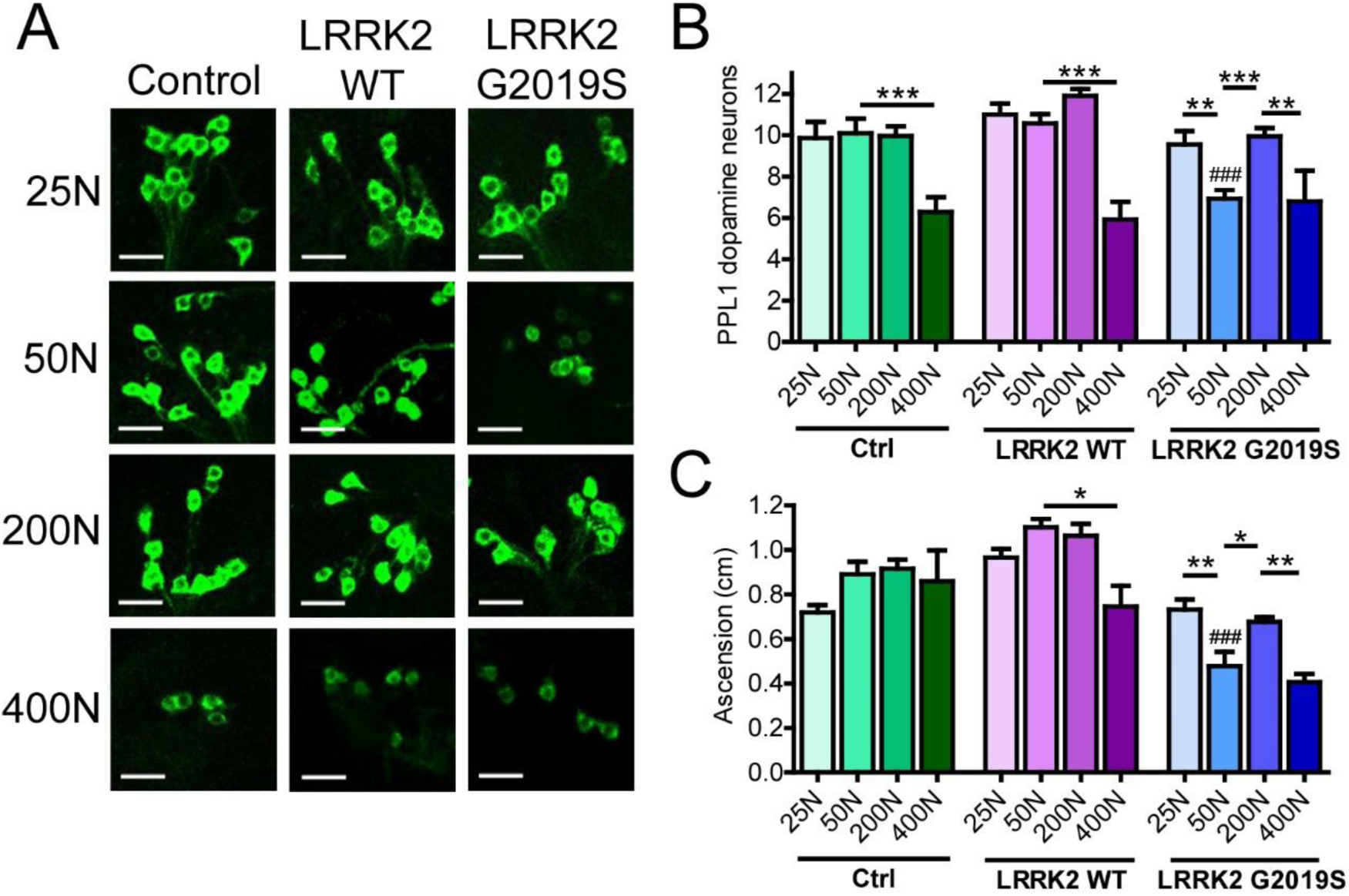
Dietary amino acids impact PD-related phenotypes in aged flies. (a,b) Effect of dietary amino acid levels on dopamine neuron number (immunostained for tyrosine hydroxylase) in the protocerebral posterior lateral 1 (PPL1) cluster. Projection images from the right hemisphere of 6-week-old Control, LRRK2 WT and LRRK2 G2019S fly brains are shown. Individual ANOVAs, Bonferroni post-tests; **p<0.01 and ***p<0.001 for diet effect and ^###^p<0.001 for genotype effect, n=12-15 brains/genotype/diet. Scale bars, 40 µm. (c) Negative geotaxis performance of 6-week-old flies (individual ANOVAs, Bonferroni post-test, *p<0.05 for diet effect, ^###^p<0.001 for genotype effect, n=4 groups of 25 flies/condition). Genotypes are Control (*Ddc-GAL4*), LRRK2 WT (*Ddc-GAL4/UAS-LRRK2 WT*) and G2019S (*Ddc-GAL4/UAS-LRRK2 G2019S*) Data are mean ± SEM.

Food intake, measured by ingestion of blue dye-labeled food, is inversely proportional to dietary amino acid concentration in 6-week-old flies (Fig. S1b), likely due to the rapid satiating effects of high amino acid diets, as has been reported for high protein diets in *Drosophila* (Sun *et al*. 2017), rodents (Solon-Biet *et al*. 2014) and humans (Westerterp-Plantenga *et al*. 2009). LRRK2 toxicity is kinase-dependent (Greggio *et al*. 2006) and to probe whether dietary amino acid concentration affects LRRK2 expression or activity in a manner that might modulate PD-related phenotypes, we assessed LRRK2 expression and LRRK2 G2019S auto-phosphorylation at Ser1292 as a readout of its kinase activity (Sheng *et al*. 2012). We find no evidence of altered LRRK2 expression or kinase activity in the heads of pan-neuronal LRRK2 transgenic flies (Fig. S1c) across dietary amino acid levels. This is further supported by the lack of dietary influence on phosphorylation of the LRRK2 substrate RPS15 (Fig. S1d).

Prior to this study, we consistently observed LRRK2 G2019S-induced dopamine neuron loss and locomotor deficits in 6-week-old flies fed standard chemically-undefined food using the same *Ddc-GAL4* and *UAS-LRRK2 G2019S* fly strains (Martin *et al*. 2014b). An estimation of the amino acid nitrogen content of the standard food used in those studies (see Methods), showed a closest match to the 50N diet used here, where we observe the same PD-related phenotypes. Collectively, these results indicate that the manifestation of LRRK2 G2019S-induced neurodegeneration in *Drosophila* is highly dependent on dietary amino acid content and, relative to food used in our previous studies which is closest to the 50N diet, can be alleviated at both lower (25N) and higher (200N) amino acid diets.

### Modulation of LRRK2 neurotoxicity by low amino acid diets is associated with attenuated protein synthesis

Since aberrant protein synthesis contributes to LRRK2 G2019S-mediated neurodegeneration in multiple models, we hypothesized that dietary amino acid levels modulate neuronal degeneration by directly impacting protein synthesis. To test this, we measured protein synthesis in aged pan-neuronal LRRK2 G2019S-expressing fly brains *ex vivo*. As expected, there is a trend towards elevated ^35^S-methionine/cysteine labeling of newly-synthesized proteins at higher dietary amino acid levels in the heads of all genotypes (Fig. 2a,b). LRRK2 G2019S flies aged on a 50N diet exhibit higher global protein synthesis relative to control genotype flies aged on the same diet (Fig. 2a,b). This is consistent with a role for elevated protein synthesis in LRRK2 G2019S-induced neurodegeneration on this diet. There is a significant reduction in protein synthesis between 50N and 25N in LRRK2 G2019S flies, suggesting that lowering protein synthesis suppresses LRRK2 G2019S neurodegeneration. This is in accordance with prior studies using protein synthesis inhibitors (Martin *et al*. 2014a; Martin *et al*. 2014b). Protein synthesis is sensitive to changes in amino acid levels in part via regulation of TOR complex 1 (TORC1) signaling (Kim *et al*. 2002). TORC1 co-ordinates nutrient amino acid input with protein synthesis output through multiple channels including direct phosphorylation of downstream targets such as S6K (ribosomal protein S6 kinase) and 4E-BP (eIF4E binding protein). Surprisingly, there is no clear effect of dietary amino acid concentration on TORC1 activity assessed via 4E-BP and S6K phosphorylation in 6-week-old fly heads (Fig. 2 c-e). While there is a trend for higher 4E-BP phosphorylation with increasing amino acids, consistent with the upregulation of protein synthesis, S6K phosphorylation is fairly constant on all diets. Interestingly, phospho-S6K is significantly higher in LRRK2 G2019S flies relative to control lines at all dietary amino acids, raising the possibility that S6K could be a LRRK2 kinase substrate. However, since S6K is not directly phosphorylated by LRRK2 in *in vitro* kinase assays (not shown), this seems to be an unlikely explanation. These results indicate that the 25N diet suppresses bulk translation in aged mutant LRRK2 flies without any prominent impact on TORC1 activity at 6 weeks.

**Figure 2.**
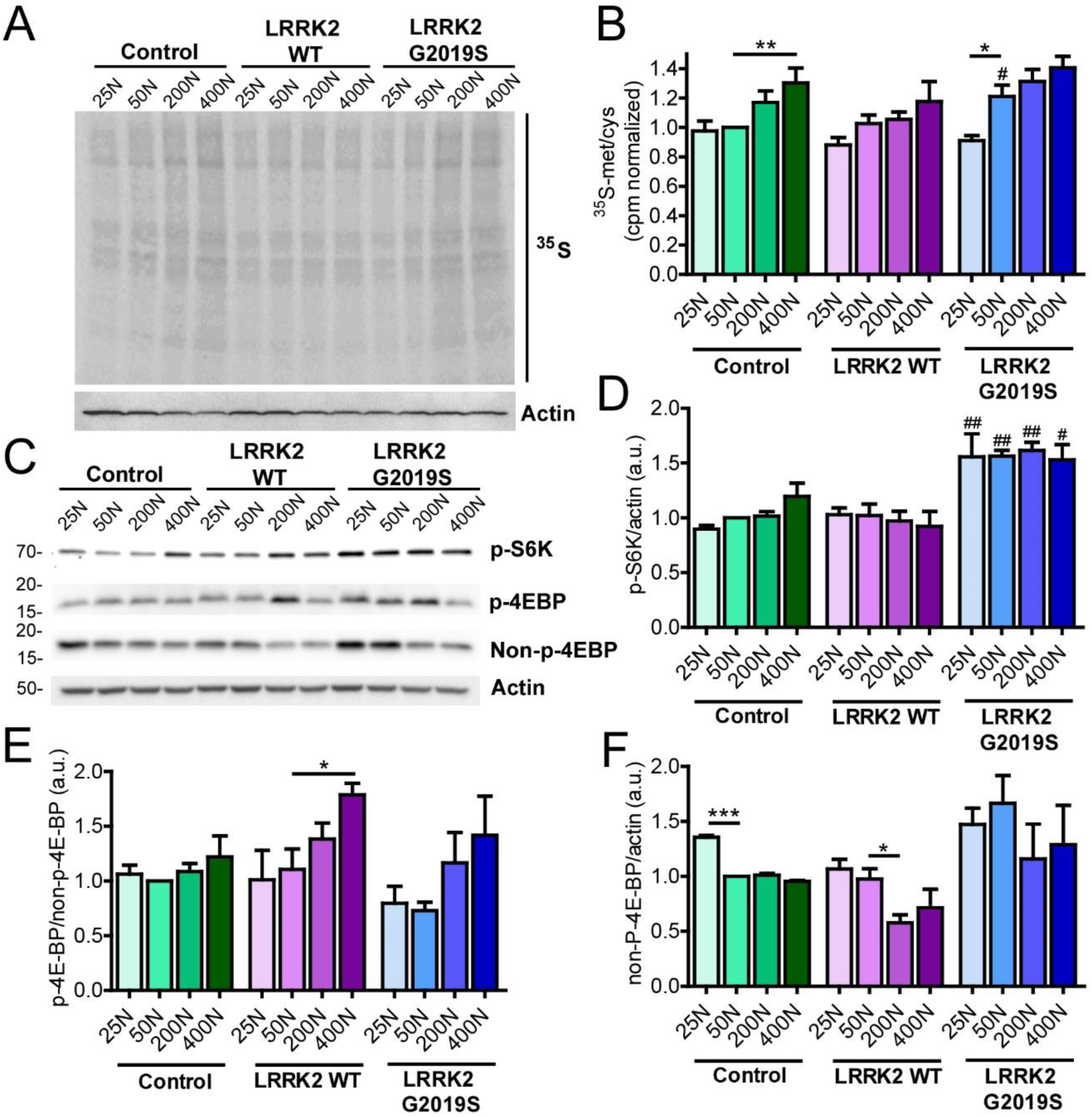
Dietary amino acids affect protein synthesis and subtly affect TORC1 activity in aged flies. (a) Representative autoradiograph for dietary amino acid effects on protein synthesis measured by ^35^S-met/cys incorporation in 6-week-old fly brains *ex vivo* (b) Radioactive CPM were divided by number of brains and normalized to Control 50N values. Individual ANOVAs, Bonferroni post-tests; *p<0.05, **p<0.01 for diet, #p<0.05 for genotype, n=6 replicates of 8 brains/genotype/condition. (c-e) TORC1 activity was measured via phospho-S6K and phospho-4E-BP levels in 6-week-old fly heads. Effect of genotype but no effect of diet on phospho-S6K (two-way ANOVA, Bonferroni post-tests ^##^p<0.01 relative to Control flies at the same diet). No significant effect of diet on phospho-4E-BP levels except for LRRK2 flies (Individual ANOVAs, Bonferroni post-test *p<0.05). (f) There is an effect of diet on non-phospho-4E-BP in Control and LRRK2 flies but not G2019S (individual ANOVAs, Bonferroni post-test ^#^p<0.05). Genotypes are Control (*elav-GAL4*), LRRK2 WT (*elav-GAL4/UAS-*LRRK2 WT) and G2019S (*elav-GAL4/UAS-LRRK2 G2019S*). Data are mean ± SEM.

Another cellular mechanism for sensing low amino acid availability is through the activation of the protein kinase GCN2 (general control nonderepressible 2) by uncharged tRNAs which promotes eIF2α phosphorylation and consequent downregulation of bulk protein synthesis (Soultoukis and Partridge 2016). However, phospho-eIF2α levels are unaffected by dietary amino acid levels within the range used here (Fig. S2), arguing against this as a possible cause for suppressed protein production. In addition to regulation by nutrient-sensing pathways, protein synthesis is also dependent on the pool of amino acids available for tRNA charging (Elf and Ehrenberg 2005). This can be readily observed in rapidly dividing organisms such as E. coli, where protein synthesis rates are clearly limited by the rate of supply of amino acids (Elf and Ehrenberg 2005). To explore whether a reduced pool of amino acids in the 25N diet accounts for the reduction in protein synthesis, we used a modified 50N diet (25N-F/H/W) in which the concentration of just three amino acids; phenylalanine (F), histidine (H), and tryptophan (W), are reduced to 25N levels while all other amino acids remain at 50N levels. These three amino acids are not thought to have a major impact on TORC1 activity, based on experiments in HEK-293T cells (D. Sabatini, personal communication), and are essential amino acids which cannot be synthesized therefore must be supplied in food. Informatively, reducing just these amino acids to 25N levels lowers protein synthesis in the heads of LRRK2 G2019S flies to virtually the same rates as those seen in flies maintained on the standard 25N diet (Fig. 3). Therefore, restricted amino acid availability in 25N fly diets appears sufficient to prevent manifestation of PD-related phenotypes. To determine whether restricting dietary amino acid content can also modulate LRRK2 toxicity in mammalian systems, we used a rat cortical neuron model of acute LRRK2 toxicity. It is well established that LRRK2 G2019S causes kinase-dependent neurotoxicity in rodent neuron cultures characterized by progressive neurite loss and cell death (Macleod *et al*. 2006). Strikingly, reducing the concentration of all amino acids in the culture medium by 20-40% during the time of LRRK2 G2019S expression ameliorates neurite loss and cell death caused by mutant LRRK2 (Figs. 4 and S3). This is accompanied by inhibition of global protein synthesis, measured via SUnSET assay (Schmidt *et al*. 2009) (Figs. 4 and S3). At a 60% reduction in amino acid concentration, neurite loss and cell death phenotypes become evident in both LRRK2 WT- and LRRK2 G2019S-transfected neurons (Figs. 4 and S3), possibly due to a starvation effect at this extent of amino acid restriction. Hence, lowering amino acid levels within a range, attenuates LRRK2 G2019S-associated neurotoxicity in both fly brains and mammalian neurons.

**Figure 3.**
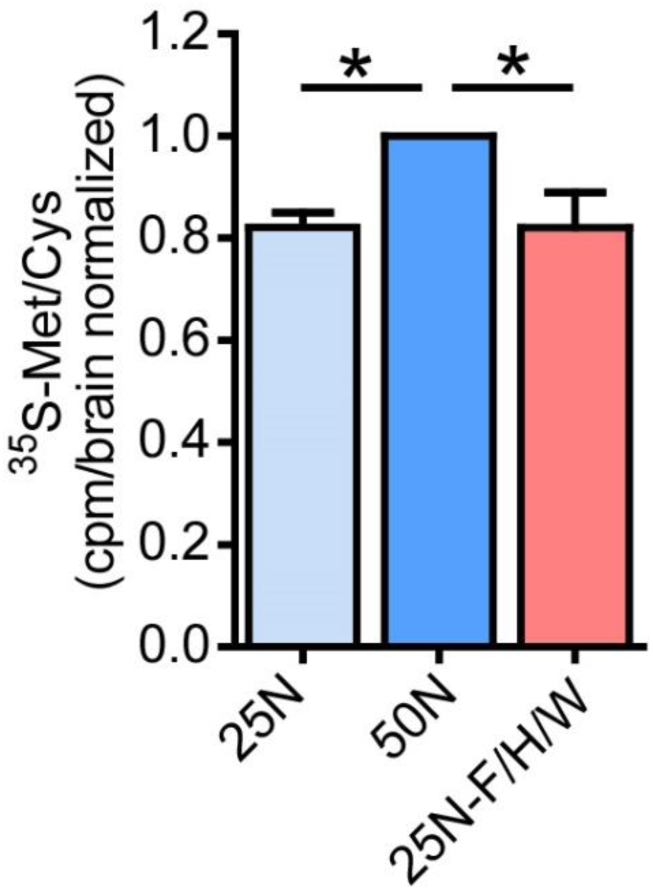
Reducing levels of Phe, His and Trp is sufficient to suppress bulk translation in aged G2019S flies. Protein synthesis was measured via 35S-met/cys incorporation in 6-week-old LRRK2 G2019S (elav-GAL4; UAS-LRRK2 G2019S) fly brains ex vivo. Protein synthesis is reduced in 25N or 25N-F/H/W (in which levels of phenylalanine (F), histidine (H) and tryptophan (W) are reduced to 25N levels) diets relative to 50N. Scintillation count values were divided by total number of brains per sample. ANOVA, Bonferroni post-test *p<0.05, n=5 replicates of 8 flies/condition. Data are mean ± SEM.

**Figure 4.**
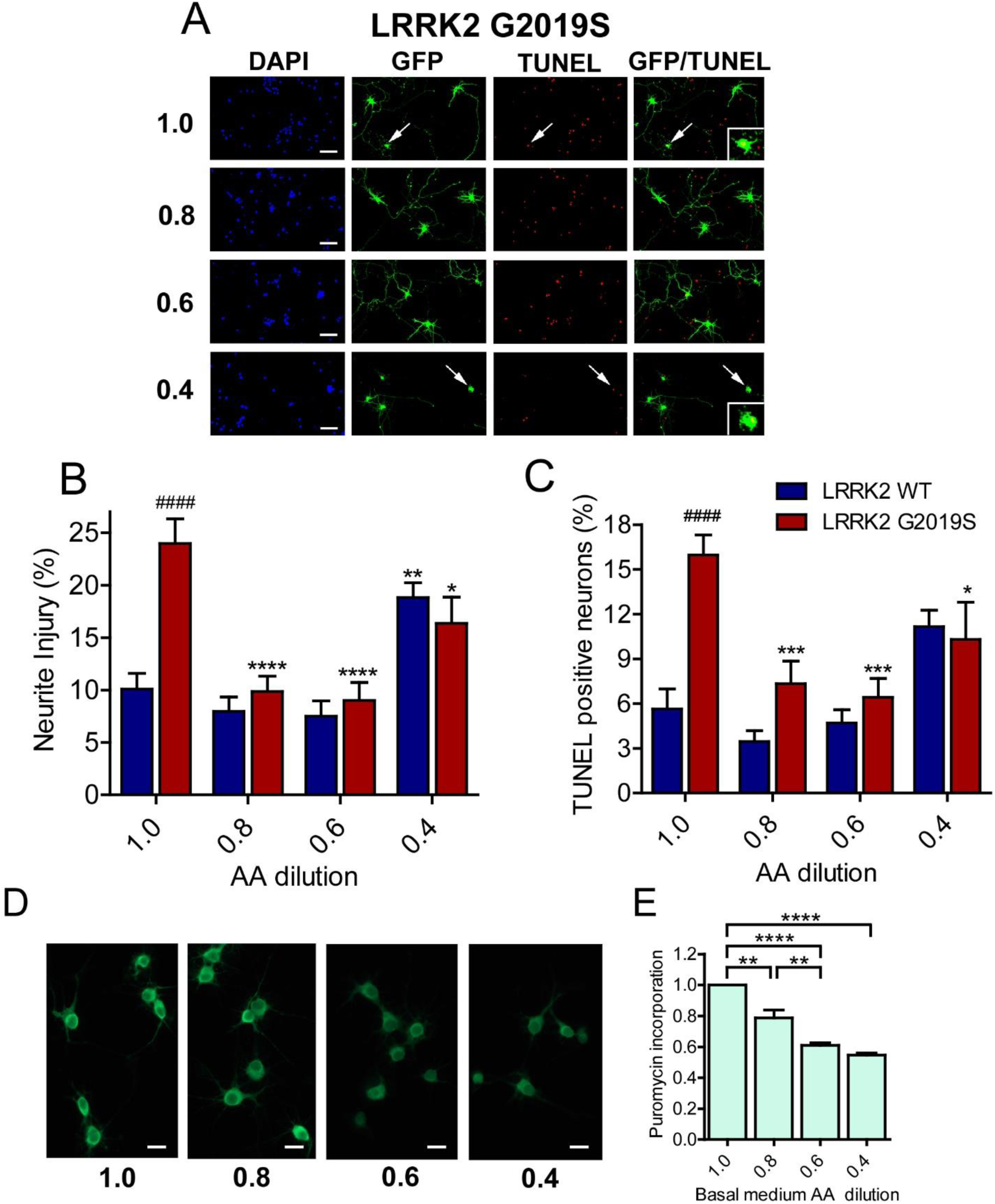
Amino acid dilution protects against LRRK2 toxicity in rat primary neurons. (a-c) Effect of culture medium amino acid dilution on LRRK2 G2019S toxicity assessed in rat cortical neurons. See Fig S3 for LRRK2 WT images. Arrows indicate neurons lacking neurites that are TUNEL-positive (see inset magnifications). Amino acid dilutions protect against LRRK2 G2019S-induced neurite injury and cell death (two-way ANOVA, Bonferroni post-test, *p<0.05, **p<0.01, ***p<0.001 relative to 1.0 condition for genotype; ###p<0.001 relative to LRRK2 control for diet, n = 6). 0.4-fold dilution lead to increased neurite injury for the LRRK2 control (Bonferroni post-test, *p<0.05). (d,e) Effect of amino acid dilution on protein synthesis assessed by puromycin incorporation. Protein synthesis is progressively reduced by a 1.0 to 0.4-fold amino acid dilution (ANOVA, Bonferroni post-test, **p<0.01, n = 3 replicates of 20 neurons/condition). Data are mean ± SEM.

### AMPK and autophagy activation on high amino acid diets block PD-related phenotypes in flies

The 200N diet effectively suppresses neurodegenerative phenotypes in aged LRRK2 G2019S flies without reducing protein synthesis or mortality (Figs. 1 and 2). This indicates that dopamine neuron viability is not always vulnerable to elevated protein synthesis. The data from S6K and 4E-BP phosphorylation suggest that there is no major effect of increasing dietary amino acid levels on TORC1 signaling in aged flies. Previous studies indicate that chronic TORC1 hyperactivity in flies can result in a stress response which triggers feedback inhibition on TORC1 signaling (Lee *et al*. 2010). To assess whether chronic consumption of high amino acid diet triggers TORC1 feedback inhibition in aged flies, we measured TORC1 activity in 1-week-old flies maintained on the same series of holidic diets for the first week of adulthood (Fig. 5). At this age, we observe a more pronounced effect of dietary amino acid levels on TORC1 activity relative to that seen in 6-week-old flies (Fig. 2), particularly at 400N. This is consistent with an age-dependent dampening of TORC1 signaling.

**Figure 5.**
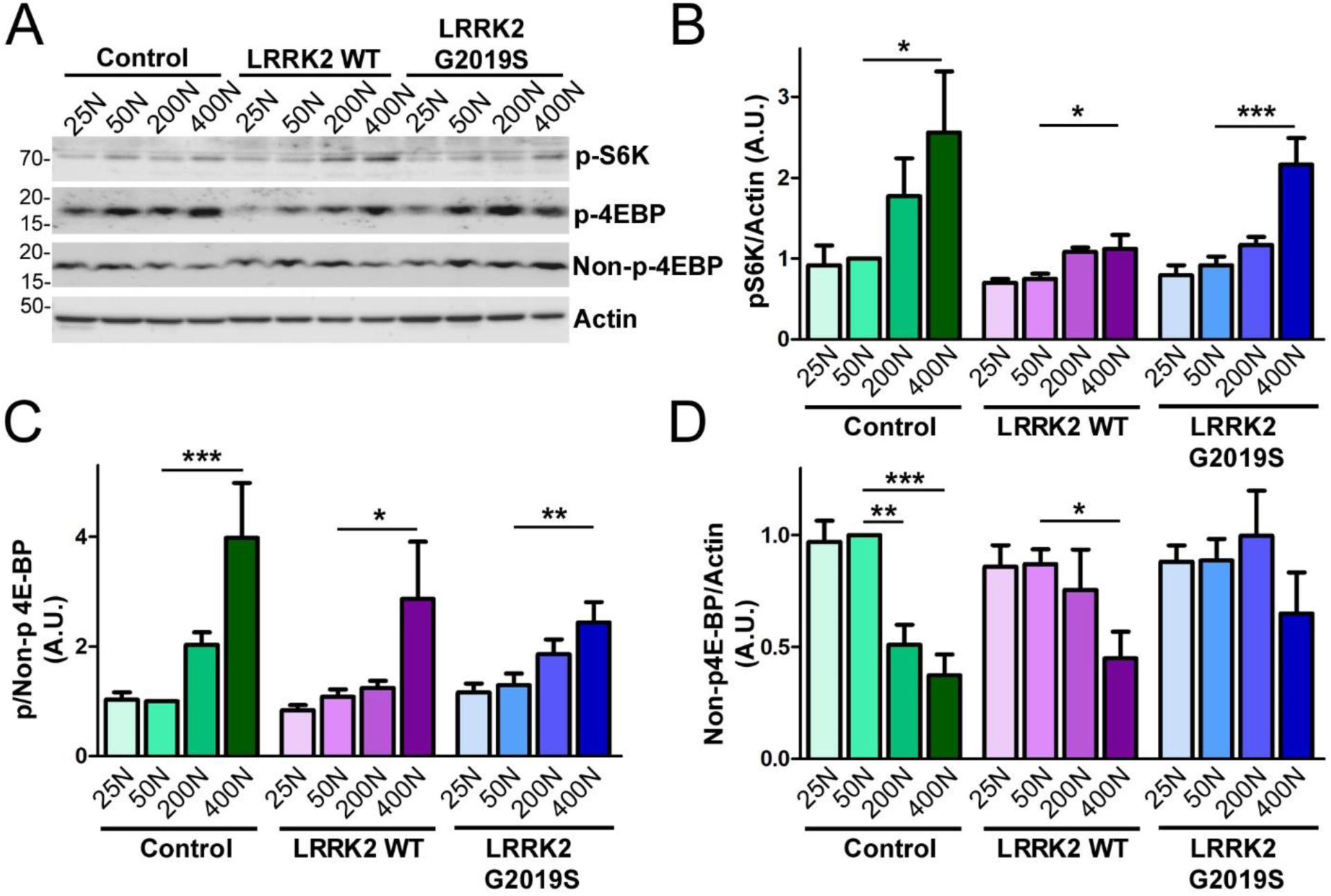
Impact of dietary amino acids on TORC1 activity in 1-week-old flies. (a-c) TORC1 activity was measured via phospho-S6K and phospho-4E-BP levels in 1-week-old fly heads. Diet affected phospho-S6K levels and ratio of phospho-4E-BP:non-phospho-4E-BP (individual ANOVAs, Bonferroni post-test, *p<0.05, **p<0.01, ***p<0.001 relative to 50N diet for genotype, n = 5). (d) Non-phospho-4E-BP levels were decreased at 200N and/or 400N diets in Control and LRRK2 flies (individual ANOVAs, Bonferroni post-test, *p<0.05, **p<0.01, ***p<0.001, n = 5). Genotypes are Control (*elav-GAL4/+*), LRRK2 WT (*elav-GAL4/*LRRK2 WT) and G2019S (*elav-GAL4/LRRK2 G2019S*). Data are mean ± SEM.

In *Drosophila*, chronically hyperactive TORC1 has been shown to induce the sestrin dSesn, which exerts feedback inhibition on TORC1 partly through direct activation of AMPK (AMP-activated protein kinase) and results in protection against age-related pathologies (Lee *et al*. 2010). Indeed, we observe that the 200N and 400N diet in LRRK2 G2019S flies and the 400N diet in control flies invokes an age-associated induction of dSesn expression at 5.5 weeks of age, which is absent in younger flies (Figs. 6a and S4). The dSesn upregulation by 200N and 400N diets is accompanied by evidence of AMPK activation (increased levels of pAMPK Thr172) in LRRK2 G2019S flies (Fig. 6b and C), which likely dampens the effects of dietary amino acids on TORC1 activity since AMPK is a known inhibitor of TORC1. As we find reduced food intake in flies maintained on higher amino acid diets (Fig. S1b), AMPK could potentially be activated in response to energy stress by sensing elevated AMP:ATP and ADP:ATP ratios. To test this possibility, we assessed ADP:ATP ratios in aged fly heads but find no significant effect of diet in any genotype, suggesting that high amino acid diets do not cause an overt deficit in ATP (Fig. 6d). To determine whether AMPK is involved in the alleviation of neurodegenerative phenotypes on the 200N diet, we silenced AMPK and assessed locomotor function in LRRK2 G2019S flies aged on 200N as well as 25N and 50N diets. RNAi-mediated knock-down of AMPK clearly blocks the rescue effect of the 200N diet (Fig. 6e), but has no significant effect on the locomotor deficit seen in LRRK2 G2019S flies aged on a 50N diet, or on locomotor abilities of LRRK2 G2019S flies aged on a 25N diet (Fig. S5a and b). Hence, AMPK is necessary for the rescue effect of the 200N diet, but not of the 25N diet. AMPK signaling activates multiple cytoprotective pathways, including autophagy. Accordingly, expression of the autophagosome components atg8a and atg9 along with the lysosomal marker LAMP1 are elevated on 200N and 400N diets (Fig. 7a-d), indicating an induction of autophagy on high amino acid diets. To determine if autophagy plays a role in suppressing neurodegenerative phenotypes on a 200N diet, we functionally silenced the autophagy-initiating complex gene atg1 (ortholog of mammalian ULK1). Previous studies have shown that autophagy cannot be induced in flies expressing a kinase-dead atg1 mutant (atg1^KQ#5B^), which functions as a dominant-negative form of this protein (Berry and Baehrecke 2007; Scott *et al*. 2007). Expression of dominant negative atg1 or RNAi-mediated atg1 silencing ablates the rescue effect of a 200N diet on the LRRK2 G2019S locomotor phenotype (Fig. 7e) suggesting that augmented autophagy contributes to this rescue. Additionally, there are no effects of atg1 loss-of-function on the LRRK2 G2019S locomotor phenotype on 50N diet, or on locomotor function of these flies on a 25N diet (Fig. S5c and d). Taken together, these data support a role for AMPK and autophagy activation in the neurodegeneration-suppressing effects of the 200N diet, but not that of the 25N diet.

**Figure 6.**
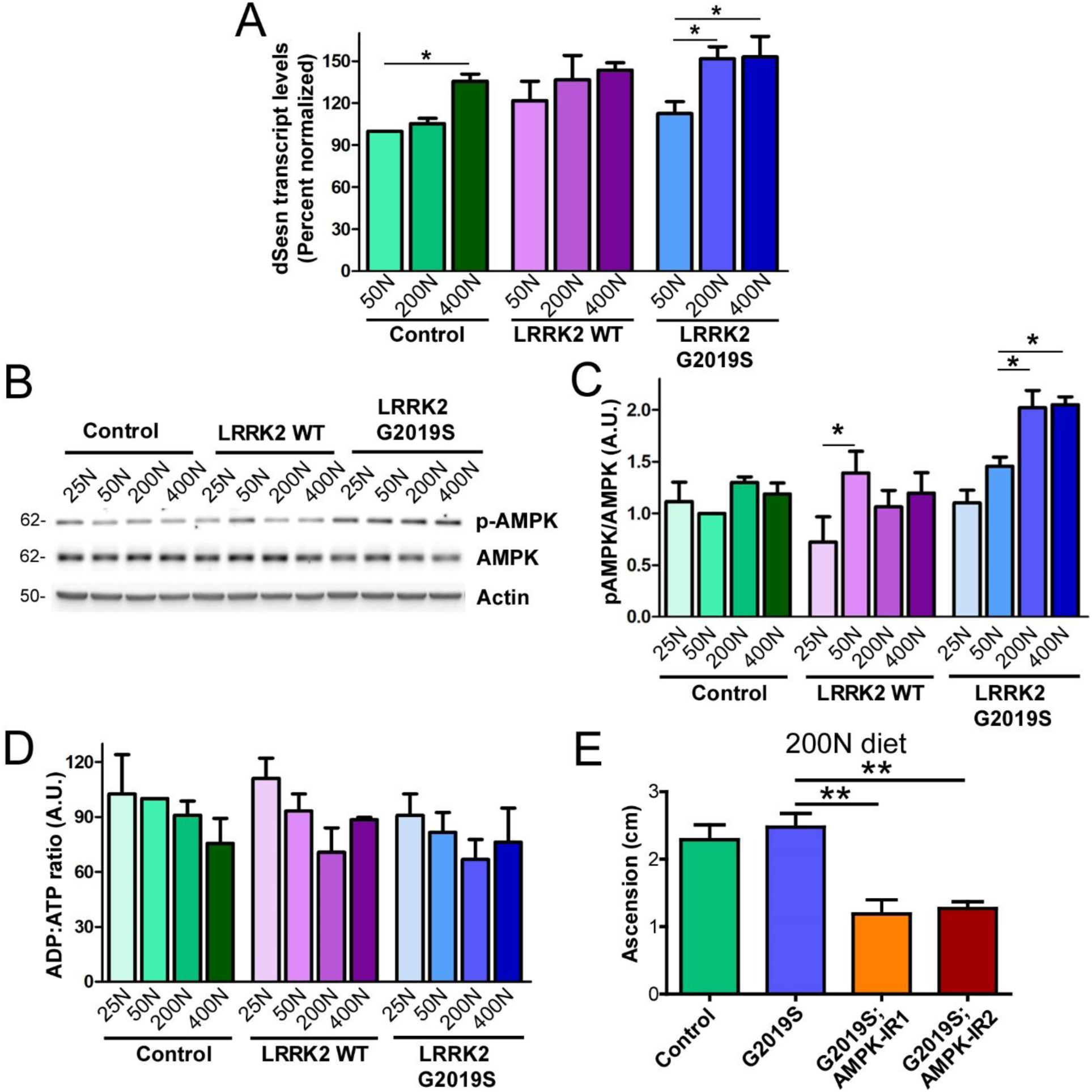
High amino acid diets induce sestrin (dSesn) expression and activate AMPK activity which is protective. (a) dSesn mRNA levels, measured by qPCR from 5.5-week-old fly heads were increased at 200N and/or 400N diets (individual ANOVAs, *p<0.05, n=3; 15 flies/condition). (b,c) AMPK activation, assessed via Thr172 phosphorylation in 6-week-old fly heads (n=3; 25 flies/condition), was increased across dietary amino acid elevation for LRRK2 and G2019S (individual ANOVAs, Bonferroni post-test *p<0.05). (d) no significant effect of dietary amino acids on ADP:ATP ratio in 6-week-old fly heads (e) AMPK knock-down via RNAi blocked the rescue effect of 200N diet on locomotor function in 6-week-old flies (ANOVA, Bonferroni post-test, **p<0.01, n = 4 groups of 25 flies/condition). Genotypes are (a-d) Control (*elav-GAL4*), LRRK2 WT (*elav-GAL4; UAS-LRRK2 WT*) and G2019S (*elav-GAL4; UAS-LRRK2 G2019S*) or (e) Control (*Ddc-GAL4*), LRRK2 WT (*Ddc-GAL4; UAS-LRRK2 WT*) and G2019S (*Ddc-GAL4; UAS-LRRK2 G2019S*). Data are mean ± SEM.

**Figure 7.**
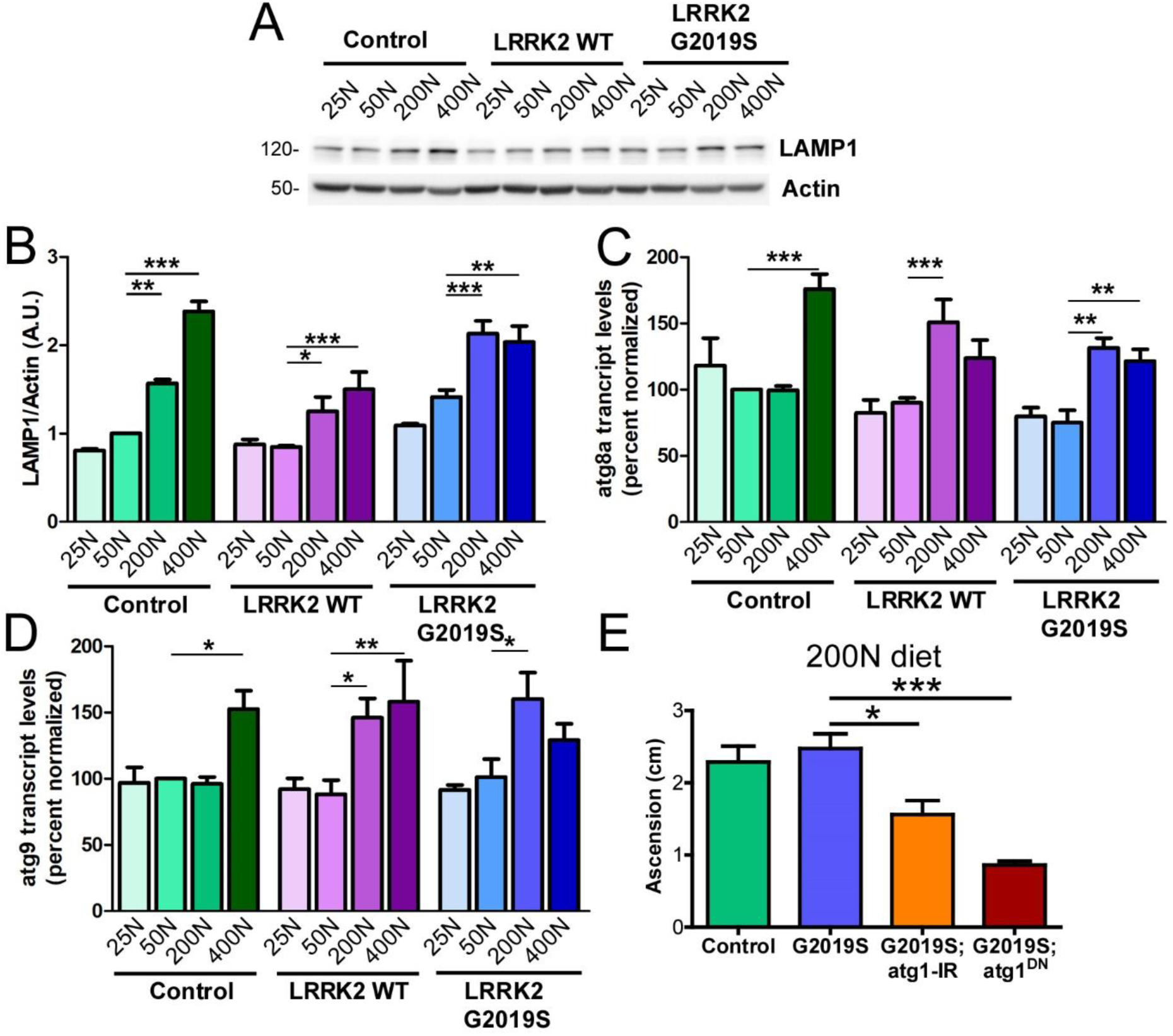
Autophagy plays a critical role in 200N diet-associated protection. (a,b) Elevated LAMP1 levels across increased dietary amino acids in 6-week-old fly heads (individual ANOVAs, Bonferroni post-tests *p<0.05, **p<0.01, ***p<0.001, n=3 groups of 25 flies/condition). Autophagosome components atg8a (c) and atg9 (d) increased transcript levels at 200N and/or 400N diets (individual ANOVAs, Bonferroni post-tests, *p<0.05, **p<0.01, ***p<0.001, n =5 groups of 15 flies/condition). (e) atg1 knock-down via RNAi (atg1-IR) or dominant-negative expression (atg1^DN^) blocked the rescue effect of 200N diet on locomotor function in 6-week-old flies (ANOVA, Bonferroni post-test, **p<0.01, n = 4 groups of 25 flies/condition). Genotypes are (a-d) Control (*elav-GAL4*), LRRK2 WT (*elav-GAL4; UAS-LRRK2 WT*) and G2019S (*elav-GAL4; UAS-LRRK2 G2019S*) or (e) Control (*Ddc-GAL4*), LRRK2 WT (*Ddc-GAL4; UAS-LRRK2 WT*) and G2019S (*Ddc-GAL4; UAS-LRRK2 G2019S*). Data are mean ± SEM.

### Pro-apoptotic markers are elevated on a 400N amino acid diet independent of LRRK2

The 400N diet causes PD-related phenotypes in controls as well as LRRK2 G2019S flies (Fig. 1), despite upregulated autophagy markers on this diet (Fig. 7a-d). To probe whether the observed neuronal loss in flies aged on a 400N diet is specific to dopamine neurons or reflects general brain and neuronal demise induced by this diet, we assessed pro-apoptotic markers across dietary amino acids. In *Drosophila*, apoptotic stimuli increase the expression of pro-apoptotic proteins such as Grim, Reaper and Hid which bind to dIAPs (*Drosophila* inhibitor of apoptosis proteins) and promote caspase release (Zheng *et al*. 2010). While dopamine neuron loss occurs to a similar extent on both 50N and 400N diets in LRRK2 G2019S flies, expression of both *grim* and *reaper* are elevated exclusively in 400N diet fly heads (Fig. S6). A similar increase in expression of these pro-apoptotic genes is found in aged control flies on 400N food, suggesting that neuronal apoptosis is promoted by this diet in aged flies independent of mutant LRRK2 expression.

### Protective effects of 25N and 200N amino acid diets are upheld in late life diet-switch

PD-related phenotypes manifest in LRRK2 G2019S flies at approximately 6 weeks of age, suggesting that aging is an important driver of LRRK2-mediated neurodegeneration. It has been proposed that dietary restriction can rapidly affect mortality even when introduced late in life, apparently neutralizing the effects of diet history and decreasing the short-term risk of mortality (Mair *et al*. 2003). To interrogate whether the beneficial effects of 25N and 200N diets on LRRK2 neurodegeneration require feeding throughout adulthood or whether initiating these diets later in life but prior to the onset of neurodegenerative phenotypes can still be protective, we performed a diet-switch in 5-week-old flies. LRRK2 G2019S flies were maintained on a 50N diet until 5 weeks of age and then switched to either 25N or 200N diets, or maintained on 50N food for 1 additional week (Fig. 8). Relative to flies kept on the 50N diet for the whole 6 weeks, a switch to either 25N or 200N diet for 1 week is sufficient to prevent onset of locomotor deficits at 6 weeks (Fig. 8). These results are consistent with the existence of a time window in organismal aging in which LRRK2 neurotoxicity can be prevented through amino acid manipulation.

**Figure 8.**
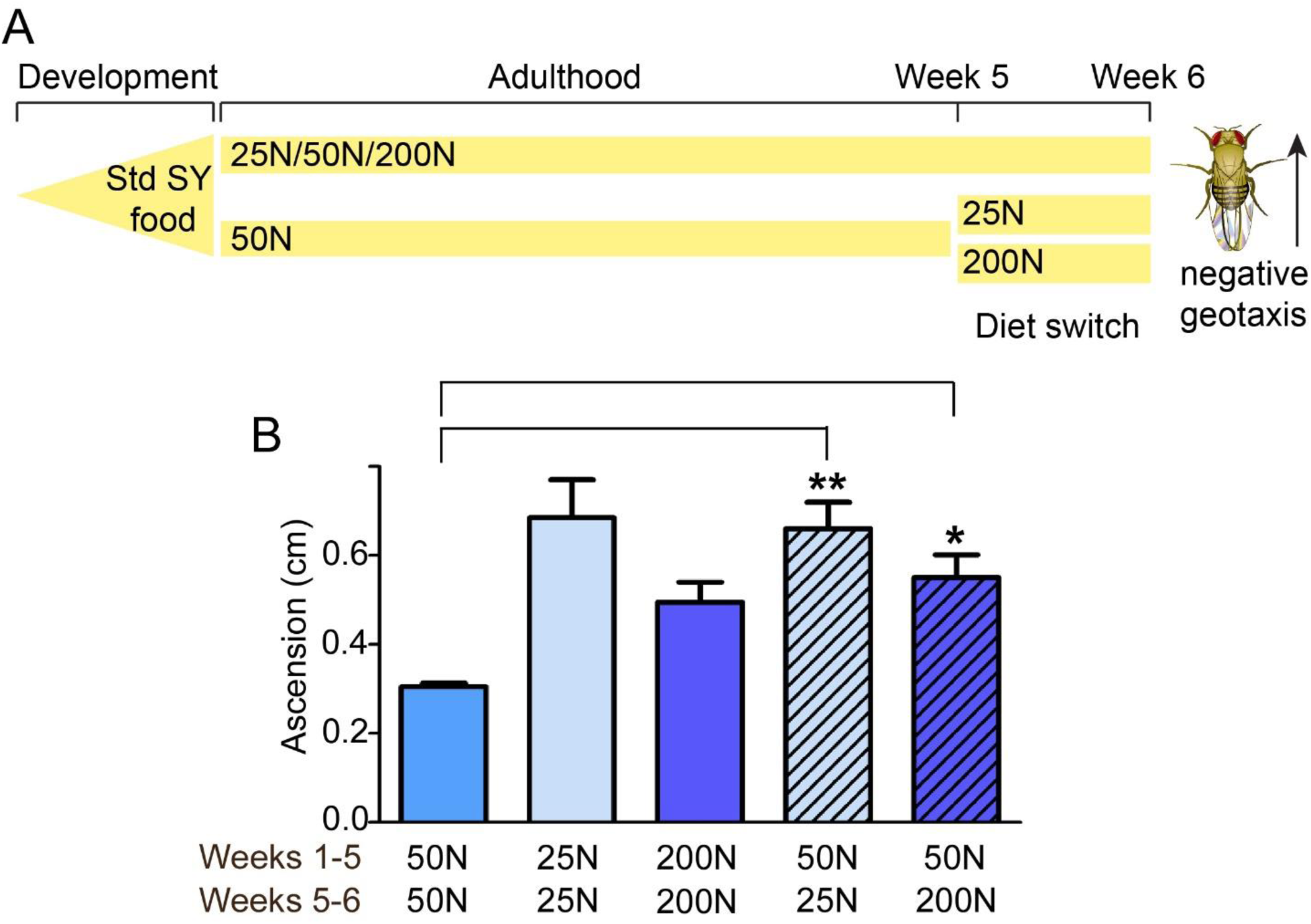
Rescue effects of dietary amino acids are retained via late life diet-switch. (a) Schematic representation of the diet switch study. A set of LRRK2 G2019S (Ddc-GAL4; UAS-LRRK2 G2019S) flies were aged on 25N, 50N or 200N diets for 6 weeks and tested for locomotor function. Another set of mutant LRRK2 flies were fed 50N diet for 5 weeks and then switched to either 25N or 200N amino acid diet for an additional 1 week (b) All flies were reared on standard sugar/yeast food. Switching flies from a 50N diet to either a 25N or 200N diet significantly improved motor function (ANOVA, Bonferroni post-test, *p<0.05, **p<0.01, n=4 groups of 25 flies/condition). Data are mean ± SEM.

## DISCUSSION

The major finding from this study is that neurodegeneration caused by the disease-associated LRRK2 G2019S mutation is highly influenced by dietary amino acid levels in adult *Drosophila* and in mammalian neuron cultures. Age-related dopamine neuron loss and motor impairment previously observed on standard chemically-undefined *Drosophila* food are recapitulated at similar amino acid levels in a holidic 50N diet fed to flies throughout adulthood. Strikingly, these phenotypes are suppressed by both restriction and moderate elevation of all dietary amino acids relative to 50N, through apparently distinct mechanisms.

In prior studies, we demonstrated that administering protein synthesis inhibitors throughout adulthood protected against elevated bulk protein synthesis and associated neurodegeneration in *Drosophila* (Martin *et al*. 2014a; Martin *et al*. 2014b). Consistent with this, restricting dietary amino acids from 50N-which is closest in amino acid content to our previous chemically-undefined food-to 25N levels similarly lowers protein synthesis and ameliorates neurodegeneration in LRRK2 G2019S flies (Figs. 1 and 2). Unexpectedly, this occurs in an apparently TORC1-independent and GCN2-independent manner (Figs. 2 and S2). As the control strains exhibit modest or no change in protein synthesis between 25N and 50N diets, a difference in dietary amino acids appears to have minimal effect on protein synthesis in control strains, while 25N levels of amino acids appear to block the LRRK2 G2019S-mediated increase in bulk protein synthesis observed at 50N and previously on chemically-undefined diets. Hence, when amino acids are more restricted such as on the 25N diet, it is possible that their levels are limiting to the point where upregulated protein synthesis, e.g through modulation of ribosomal function as proposed for LRRK2 G2019S (Martin *et al*. 2014a; Martin *et al*. 2014b) becomes inhibited. This could conceivably account for a greater impact of the 25N diet on protein synthesis in this strain. Informatively, lowered protein synthesis can be also be achieved via restricted availability of just phenylalanine, histidine and tryptophan (Fig. 3), which are not recognized TOR activators, suggesting that it is mediated through a depletion of the amino acid pool available for protein synthesis. Similarly, lowering amino acid levels in the culture medium of primary rat neurons attenuates LRRK2 G2019S-induced neurite loss and cell death with a concomitant reduction in protein synthesis (Figs. 4 and S3). Our preliminary evidence supporting a protective effect of amino acid restriction on mammalian neurons warrants future efforts to understand whether disease-related phenotypes in mammalian LRRK2 *in vivo* models can similarly be prevented.

While amino acid restriction is likely neuroprotective through lowering translation, elevating dietary amino acid concentration from 50N to 200N also blocks dopamine neuron loss and motor deficits in LRRK2 G2019S flies without lowering protein synthesis (Figs. 1 and 2). Assessment of 4E-BP and S6K phosphorylation revealed that TORC1 activity generally increases with higher dietary amino acid levels in young adult flies but that this effect becomes more modest in aged flies (Figs. 2c-e and 5a-c). With increasing levels of 4E-BP phosphorylation on higher amino acid diets, non-phosphorylated 4E-BP levels trend downward as would be expected with a higher proportion of phosphorylated protein (Figs. 2e,f and 5c,d). Sestrins, which are evolutionarily conserved stress-inducible proteins, are negative feedback regulators of TORC1 signaling and accumulate upon chronic TORC1 activation in *Drosophila* to protect against age-related pathologies (Lee *et al*. 2010). Here, we observe a transcriptional induction of the *Drosophila* sestrin dSesn, in aged flies housed on high amino acid diets (200N and/or 400N), but no effect of diet on dSesn expression in younger flies, consistent with an age-related dampening of TORC1 responsiveness to dietary amino acids (Figs. 6a and S4). As in mammals, dSesn in flies promotes activity of AMPK which in turn inhibits TORC1 signaling (Lee *et al*. 2010; Kim and Lee 2015). The strongest dSesn induction across diet occurs in LRRK2 G2019S flies, where dSesn is upregulated at both 200N and 400N diets relative to 50N, while it only appears to be elevated at 400N in controls. Interestingly, we also observe a more pronounced upregulation of AMPK activity and the autophagy marker LAMP1 in LRRK2 G2019S flies on the 200N diet than in control strains (Figs. 6b,c and 7a-d). This raises the possibility that additional stressor(s) may be present to induce dSesn expression specifically in LRRK2 G2019S flies but not controls on the 200N diet. Prior studies indicate that chronic TORC1 activity induces dSesn expression in a mechanism involving reactive oxygen species (ROS) which are also established activators of mammalian sestrins (Lee *et al*. 2010), raising the possibility that basal ROS levels might be higher in LRRK2 G2019S flies, as might be suggested by their heightened sensitivity to ROS-generating compounds such as rotenone (Ng *et al*. 2009).

An established relationship between autophagy and amino acid levels has been described at the level of amino acid starvation, which triggers autophagy induction through GCN2 activation and TORC1 silencing. We see no evidence for the GCN2 pathway being affected by the amino acid range tested here (Fig. S2), or evidence that autophagy markers are elevated by low dietary amino acid levels in this study (Fig. 7a-d). This indicates that amino acid levels on the 25N diet are not low enough to elicit a starvation response which might potentially be invoked at lower amino acid levels. The induction of autophagy on high amino acid diets associated with elevated dSesn expression and AMPK activation seen here appears to be a distinct mechanism by which amino acids impinge on autophagy. Silencing AMPK or the autophagy-initiating gene atg1 eliminates the 200N diet protective effect on LRRK2 G2019S flies, but does not impact the protective effect of the 25N diet, where AMPK and autophagy activity appear similar to those on a 50N diet (Figs. 6, 7 and S5). An attractive explanation for these findings is that AMPK activation engenders a neuroprotective effect involving autophagy that staves off neurodegeneration. Upon activation, AMPK limits energy-consuming anabolic processes while promoting catabolic processes such as autophagy.

A boost in protein turnover through autophagy could compensate for the elevated bulk protein synthesis caused by LRRK2 G2019S, thereby re-establishing protein homeostasis. Dietary promotion of autophagy via AMPK activation may also protect against the reported perturbation of lysosomal function by mutant LRRK2 (Wallings *et al*. 2019) additionally promoting a restoration of protein homeostasis. The sestrin-AMPK-autophagy axis acts to limit damage in the face of numerous cell stressors, and was previously shown to protect rodent dopaminergic cells against rotenone toxicity (Hou *et al*. 2015). Likewise, expression of constitutively-active AMPK was shown to prevent dopamine neuron loss and motor impairments in rats overexpressing alpha-synuclein (Bobela *et al*. 2017) and to protect against the same phenotypes in flies expressing LRRK2 G2019S (Ng *et al*. 2012). Additionally, it is notable that both translation suppression and AMPK activation were shown to protect against age-related pathology in parkin null flies (Tain *et al*. 2009; Ng *et al*. 2012), thus it is tempting to speculate that the beneficial effects of dietary manipulations we observe could extend beyond mutant LRRK2. Together, these findings support the potential for AMPK activation to protect against the development of PD phenotypes. Since AMPK can also be activated under conditions of energy deficit and we observe an inverse relationship between dietary amino acid levels and food intake, we considered the possibility that AMPK activation on high amino acid diets could be triggered by low ATP levels. However, ADP:ATP ratios are not affected by the diets used in the study (Fig. 6d), and additionally, egg laying was also shown in previous studies to track positively with amino acid levels (Piper *et al*. 2014) further arguing against major energy deficits which are known to impede fecundity in *Drosophila* (Partridge *et al*. 2005).

We chose to focus on manipulating dietary amino acid levels because anabolic protein synthesis is strongly linked to dietary protein levels and amino acid nutrient status. In contrast to modifying levels of whole proteins, manipulating free amino acid levels avoids the potential variability of digestion and oligopeptide transport-based uptake of individual proteins (Soultoukis and Partridge 2016) and is therefore more suitable for examining the post-absorptive effects of dietary amino acids on neurodegeneration. Additionally, evidence from animal models consistently indicates that dietary protein or free amino acid content is a major determinant of organismal life span (Grandison *et al*. 2009) invoking its potential to impact aging and age-related pathologies. Modifying dietary amino acid levels does impact food intake, resulting in an apparent satiating effect of high amino acid diets on appetite. This makes it challenging to control for intake of other dietary components across the amino acid diet range which could possibly influence the phenotypes assessed here. Importantly, we do not find any evidence of energy stress on high amino acid diet that could promote degenerative phenotypes. Future detailed investigation is warranted into the effect of manipulating other dietary components either alone or in combination, and while we did not directly explore this possibility within the current study, using the holidic diet approach should facilitate such investigations. The holidic diet was previously validated for use in adult flies to obtain optimal life span and fecundity while fly development was reported to be slightly delayed on this media (Piper *et al*. 2014). Hence, in our study, flies were reared on standard sugar, yeast food and holidic diet exposure and amino acid variation was limited to adulthood in order to avoid potential effects on development.

In summary, we find that manipulating dietary amino acid levels can potently modulate mutant LRRK2 toxicity in both *Drosophila* and mammalian systems. Our findings underscore the importance of diet in determining vulnerability of aging organisms to neurodegeneration and corroborate reports from mouse and primate studies that dietary restriction ameliorates PD-related phenotypes (Bayliss *et al*. 2016). A growing body of evidence implicates loss of translation control as a causative factor in several neurological diseases including PD, highlighting the necessity for protein homeostasis to maintain neuronal function. Further understanding the impact of dietary influences on translation in the context of these diseases will help evaluate the potential for dietary intervention to halt their onset or delay their progression.

## Acknowledgements

This work was supported by National Institutes of Health (P30NS061800) to the OHSU Advanced Light Microscopy Core, an American Parkinson Disease Association Post-Doctoral Fellowship to V.C-V. and by National Institutes of Health (K01AG050718), American Parkinson Disease Association Research Grant and OHSU Neurology Foundation Funds to I.M.

## SUPPLEMENTARY FIGURES

**Figure S1.**
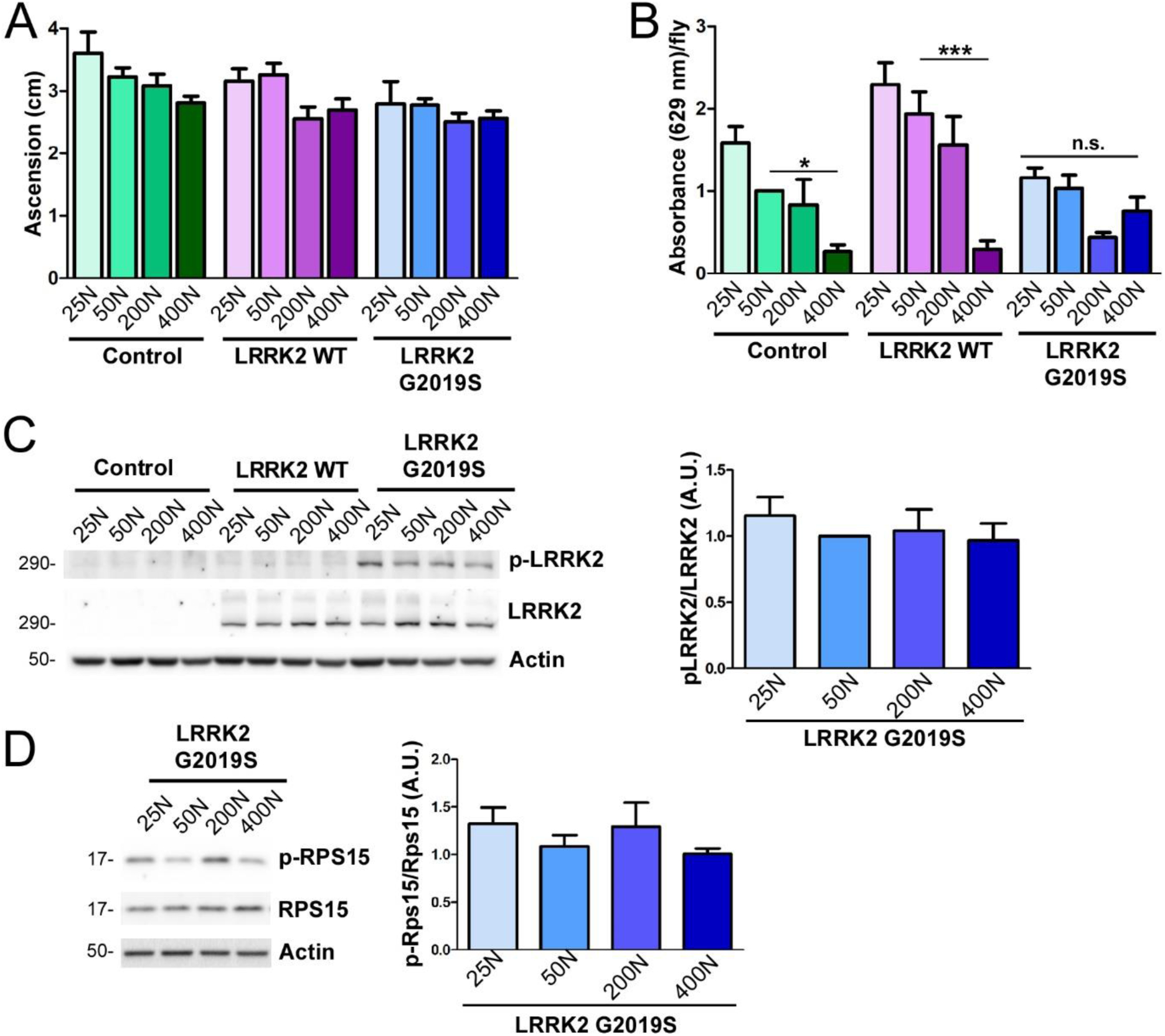
Characterization of holidic diet effects on fly locomotion, food intake and LRRK2 kinase activity. (a) No significant effect of diet on locomotor function in 1-week-old Control, LRRK2 WT or LRRK2 G2019S flies (individual ANOVAs, n.s. n = 4 groups of 25 flies/condition). Genotypes are Control (*Ddc-GAL4*), LRRK2 WT (*Ddc-GAL4; UAS-LRRK2 WT*) and LRRK2 G2019S (*Ddc-GAL4; UAS-LRRK2 G2019S*). (b) Dietary amino acids significantly impacted consumption of food for Control and LRRK2 flies but not G2019S (individual ANOVAs, Bonferroni post-test, *p<0.05, ***p<0.001, n = 3 groups of 10 flies/condition). No effect of diet on human LRRK2 kinase activity assessed by (c) auto-phosphorylation at Ser1292 relative to total LRRK2 levels in 6-week-old fly heads (ANOVA, n.s. n = 4 replicates of 25 heads/condition) or (d) phosphorylation of ribosomal protein S15 at T136 (ANOVA, n.s. n = 4 replicates of 25 flies/condition). Human LRRK2 but not *Drosophila* LRRK was immunoblotted. Genotypes in (b-d) are Control (*elav-GAL4*), LRRK2 WT (*elav-GAL4; UAS-LRRK2 WT*) and G2019S (*elav-GAL4; UAS-LRRK2 G2019S*). Data are mean ± SEM.

**Figure S2.**
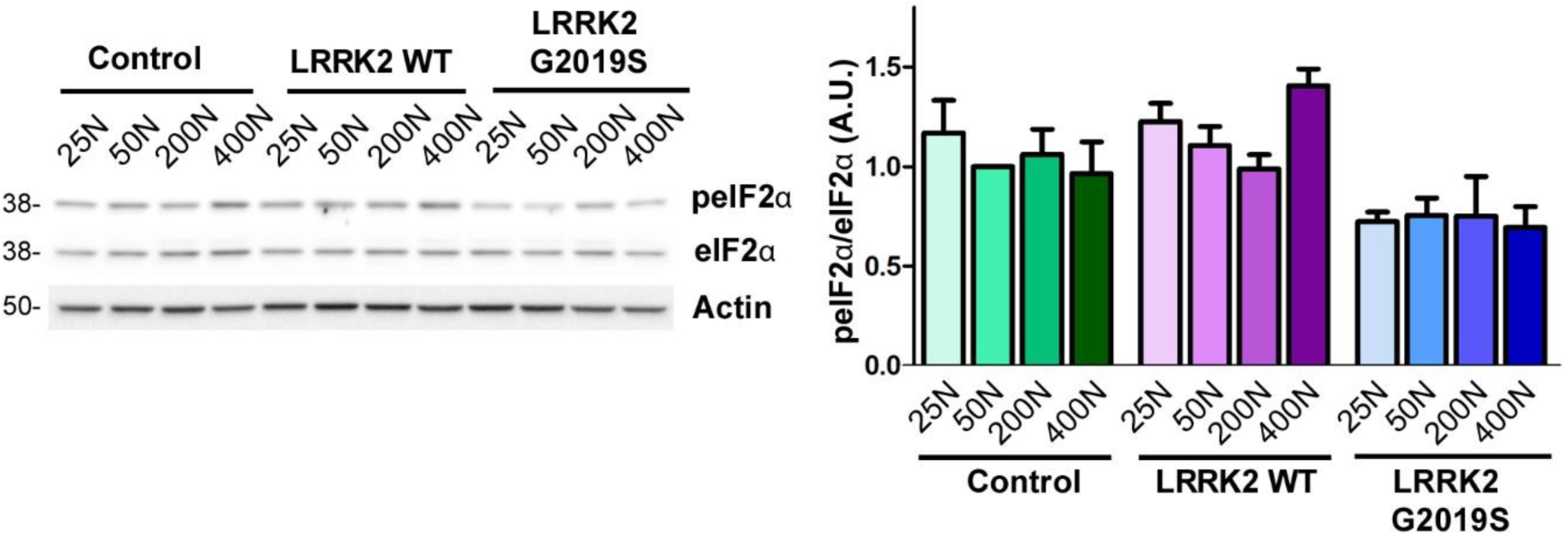
No substantive effect of dietary amino acid effects on the protein synthesis regulator eIF2α. (a) No effect of diet on GCN2 pathway activity assessed by eIF2α phosphorylation measurement (individual ANOVAs, n.s. n = 3 groups of 25 flies/condition). Genotypes are Control (*elav-GAL4*), LRRK2 WT (*elav-GAL4; UAS-LRRK2 WT*) and G2019S (*elav-GAL4; UAS-LRRK2 G2019S*). Data are mean ± SEM.

**Figure S3.**
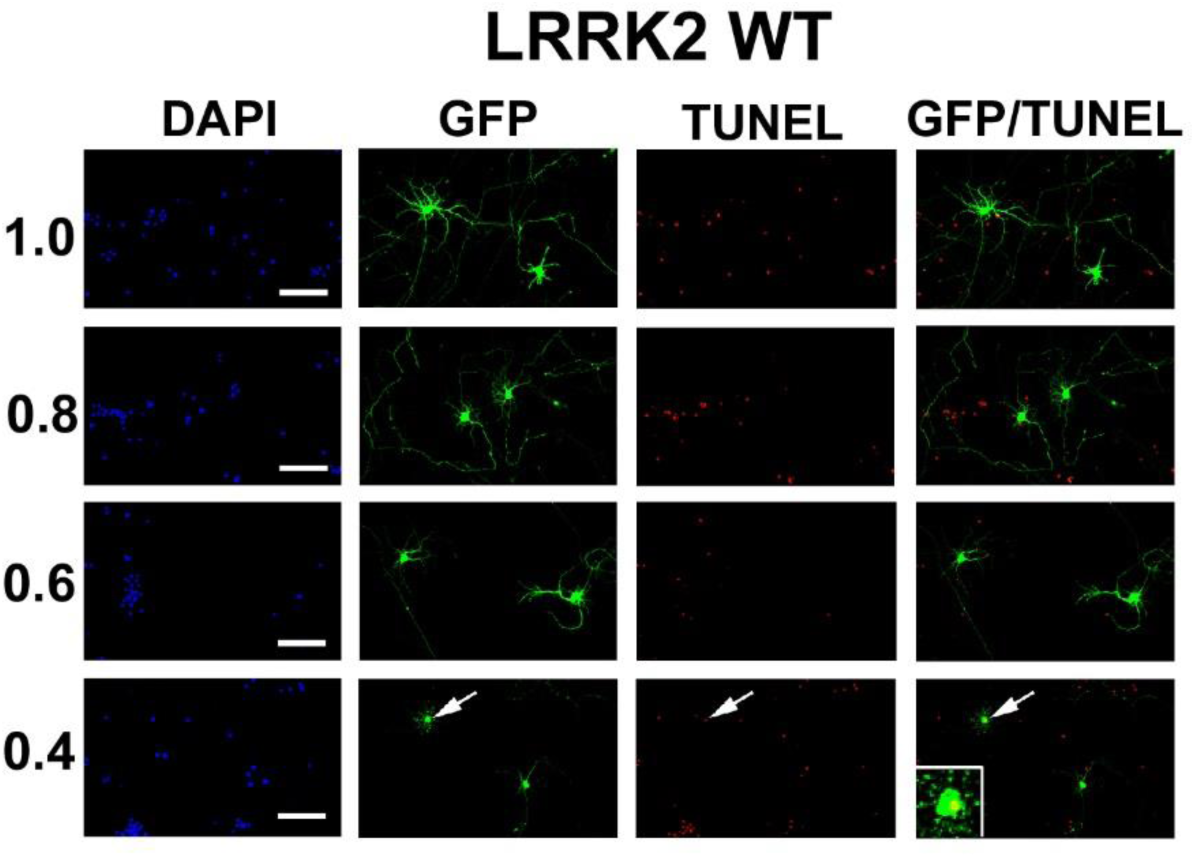
Effect of amino acid dilution on LRRK2 WT-transfected rat neurons. Images of LRRK2 WT-overexpressing neurons to accompany data in Fig. 5. Dilution of amino acids in the culture medium 0.4-fold induces toxicity (neurite loss and cell death). Arrows indicate neurons lacking neurites which are also TUNEL-positive (see inset magnifications). Scale bars, 20 µm.

**Figure S4.**
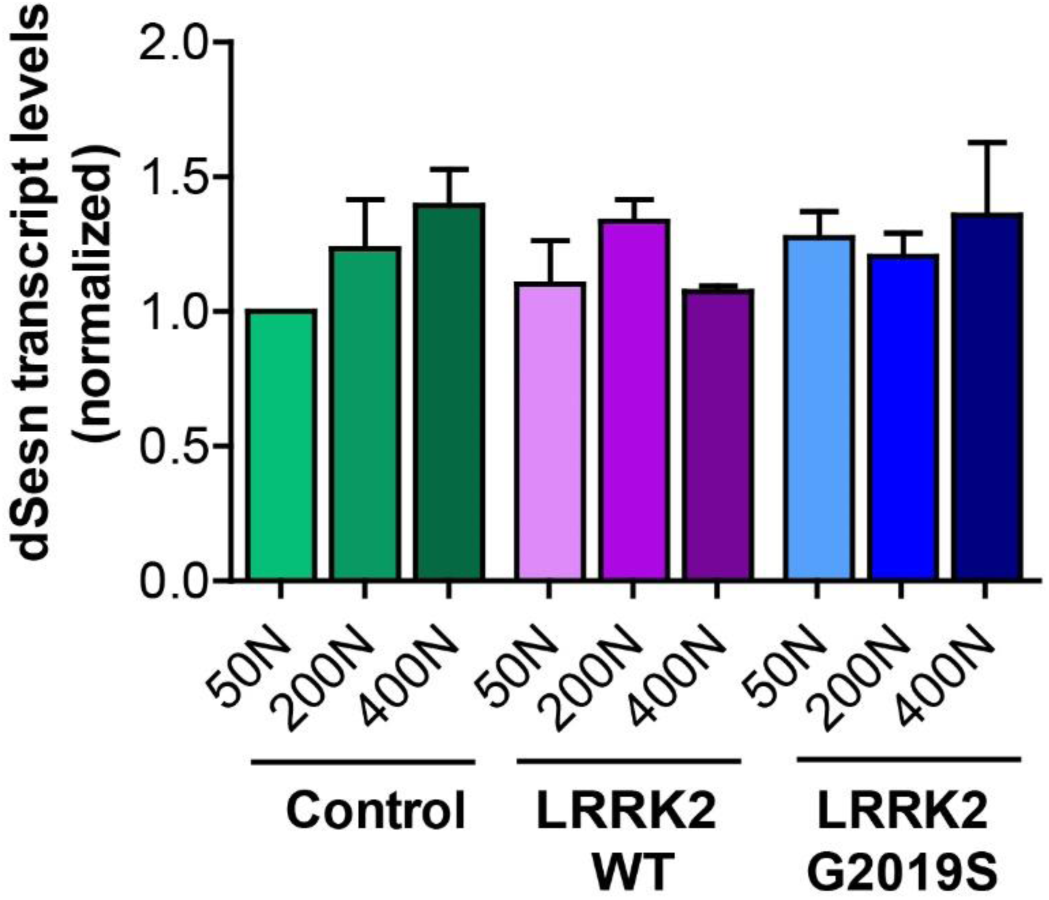
No substantive effect of diet on dSesn expression in 3-week-old flies. No significant effect of diet in Control, LRRK2 WT or LRRK2 G2019S 3-week-old fly heads assessed by qPCR (n=3 groups of 15 flies/condition). Genotypes are Control (elav-GAL4), LRRK2 WT (elav-GAL4; UAS-LRRK2 WT) and LRRK2 G2019S (elav-GAL4; UAS-LRRK2 G2019S). Data are mean ± SEM.

**Figure S5.**
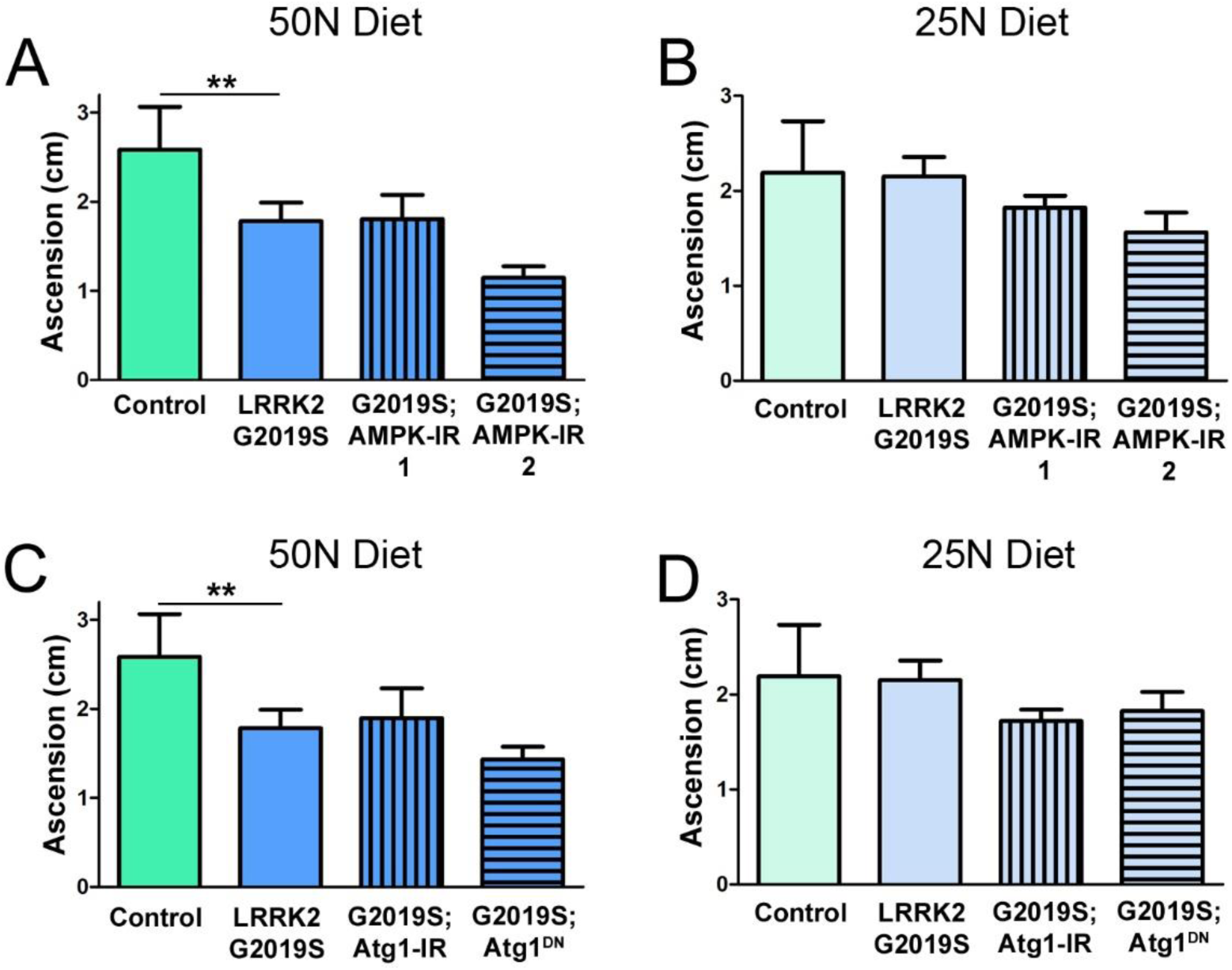
Reduced AMPK expression or atg1 expression/function do not affect motor abilities of flies aged on 50N or 25N diets. RNAi-mediated knockdown of AMPK (AMPK-IR1 or AMPK-IR2) has no significant effect on motor performance of G2019S flies on 50N (a) or 25N (b) diets for 6-weeks (individual ANOVAs, Bonferroni post-test, **p<0.01, n=4 groups of 25 flies/genotype). RNAi-mediated knock-down of atg1 or expression of dominant-negative atg1DN has no significant effect on motor performance of G2019S flies on 50N (c) or 25N (d) diets for 6-weeks (individual ANOVAs, Bonferroni post-test, **p<0.01, n=4 groups of 25 flies/genotype). Genotypes are Control (Ddc-GAL4), LRRK2 G2019S (Ddc-GAL4; UAS-LRRK2 G2019S), LRRK2 G2019S; AMPK-IR (Ddc-GAL4; UAS-LRRK2 G2019S/UAS-AMPK-IR), LRRK2 G2019S; atg1-IR (Ddc-GAL4; UAS-LRRK2 G2019S/UAS-atg1-IR) and LRRK2 G2019S; atg1DN (Ddc-GAL4; UAS-LRRK2 G2019S/UAS-atg1DN). Data are mean ± SEM.

**Figure S6.**
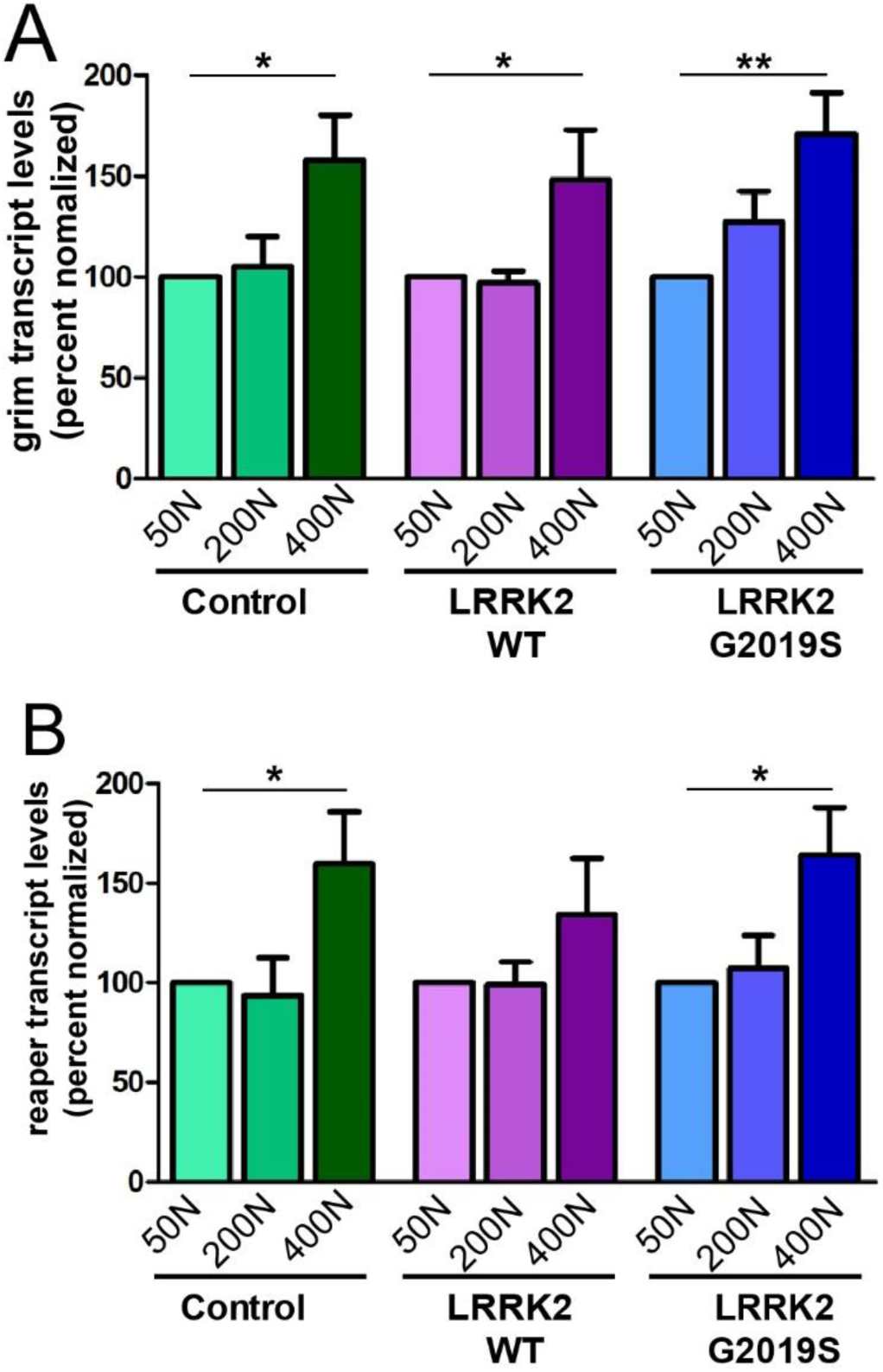
Chronic consumption of 400N diet leads to elevated pro-apoptotic gene expression. Grim (a) and reaper (b) transcript levels assessed by qPCR in 6-week old fly heads were significantly elevated on the 400N diet (individual ANOVAs, Bonferroni post-test, *p<0.05, **p<0.01, n=6; 15 flies/condition). Genotypes are Control (*elav-GAL4*), LRRK2 WT (*elav-GAL4; UAS-LRRK2 WT*) and G2019S (*elav-GAL4; UAS-LRRK2 G2019S*). Data are mean ± SEM.

## SUPPLEMENTARY TABLES

**Table S1.** Composition of holidic diet media.

| Components | Stocks | Diets |  |  |  |
| --- | --- | --- | --- | --- | --- |
| Before autoclaving |  | 25N | 50N | 200N | 400N |
| Agar (g) | - | 20 | 20 | 20 | 20 |
| L- Isoleucine (g) | - | 0.2275 | 0.455 | 1.82 | 3.64 |
| L- Leucine (g) | - | 0.1512 | 0.3025 | 1.21 | 2.42 |
| L- Tyrosine (g) | - | 0.0525 | 0.105 | 0.42 | 0.84 |
| Sucrose (g) | - | 17.12 | 17.12 | 17.12 | 17.12 |
| Base Buffer (ml)* | 10x | 100 | 100 | 100 | 100 |
| Calcium Chloride (ml) | 250 g/L | 1 | 1 | 1 | 1 |
| Magnesium Sulphate (ml) | 250 g/L | 1 | 1 | 1 | 1 |
| Copper Sulphate (ml) | 2.5 g/L | 1 | 1 | 1 | 1 |
| Iron Sulphate (ml) | 25 g/L | 1 | 1 | 1 | 1 |
| Manganese Chloride (ml) | 1 g/L | 1 | 1 | 1 | 1 |
| Zinc Sulphate (ml) | 25 g/L | 1 | 1 | 1 | 1 |
| Cholesterol (ml)* | 20 mg/mL | 15 | 15 | 15 | 15 |
| After autoclaving |  |  |  |  |  |
| Nucleic acids/Lipids (ml)* | 125x | 8 | 8 | 8 | 8 |
| Essential amino acids (ml)* |  | 7.564 | 15.128 | 60.512 | 121.02 |
| Non-essential amino acids (ml)* |  | 7.564 | 15.128 | 60.512 | 121.02 |
| Sodium Glutamate (ml) | 100 g/L | 1.89 | 3.78 | 15.12 | 30.24 |
| Vitamin (ml)* | 47.6x | 21 | 21 | 21 | 21 |
| Folic Acid (ml) | 0.5 g/L | 1 | 1 | 1 | 1 |
| Propionic Acid (ml) | - | 6 | 6 | 6 | 6 |
| Nipagin (ml)* | 100 g/L | 15 | 15 | 15 | 15 |

\* Refer to Piper et al., 2014 for complete composition

**Table S2.** Composition of Neurobasal-based medium minus phenol red and amino acids.

| **Components** | **Concentration**<br>**(mg/L)** | **mM** |
| --- | --- | --- |
| **Vitamins** |  |  |
| Choline chloride | 4 | 0.02857143 |
| D-Calcium pantothenate | 4 | 0.00838574 |
| Folic Acid | 4 | 0.0090703 |
| Niacinamide | 4 | 0.03278688 |
| Pyridoxal hydrochloride | 4 | 0.01960784 |
| Riboflavin | 0.4 | 0.00106383 |
| Thiamine hydrochloride | 4 | 0.01186944 |
| Vitamin B12 | 0.0068 | 5.02E-06 |
| i-Inositol | 7.2 | 0.04 |
| **Inorganic Salts** |  |  |
| Calcium Chloride (CaCl<sub>2</sub>)<br>(anhyd.) | 200 | 1.8018018 |
| Ferric Nitrate<br>(Fe(NO<sub>3</sub>)<sub>3</sub>·9H<sub>2</sub>O) | 0.1 | 2.48E-04 |
| Magnesium Chloride<br>(anhydrous) | 77.3 | 0.8136842 |
| Potassium Chloride (KCl) | 400 | 5.3333335 |
| Sodium Bicarbonate<br>(NaHCO<sub>3</sub>) | 2200 | 26.190475 |
| Sodium Chloride (NaCl) | 3000 | 51.724136 |
| Sodium Phosphate monobasic<br>(NaH<sub>2</sub>PO<sub>4</sub>·H<sub>2</sub>O) | 125 | 0.9057971 |
| Zinc sulfate (ZnSO<sub>4</sub>·7H<sub>2</sub>O) | 0.194 | 6.74E-04 |
| **Other components** |  |  |
| D-Glucose (Dextrose) | 4500 | 25 |
| HEPES | 2600 | 10.92437 |
| Phenol Red | 8.1 | 0.02151966 |
| Sodium Pyruvate | 25 | 0.22727273 |

**Table S3.** Composition of 25N-F/H/W holidic medium.

| Components | Stocks | Diets |
| --- | --- | --- |
| **Before autoclaving** |  | **25N-F/H/W** |
| Agar (g) | - | 20 |
| L- Isoleucine (g) | - | 0.455 |
| L- Leucine (g) | - | 0.3025 |
| L- Tyrosine (g) | - | 0.105 |
| Sucrose (g) | - | 17.12 |
| Base Buffer (ml)* | 10x | 100 |
| Calcium Chloride (ml) | 250 g/L | 1 |
| Magnesium Sulphate (ml) | 250 g/L | 1 |
| Copper Sulphate (ml) | 2.5 g/L | 1 |
| Iron Sulphate (ml) | 25 g/L | 1 |
| Manganese Chloride (ml) | 1 g/L | 1 |
| Zinc Sulphate (ml) | 25 g/L | 1 |
| Cholesterol (ml)* | 20 mg/mL | 15 |
| **After autoclaving** |  |  |
| Nucleic acids/Lipids (ml)* | 125x | 8 |
| Essential amino acids (ml)*<br>(excluding Phe, His, Trp) |  | 15.128 |
| Phe, His, Trp solution |  | 7.564 |
| Non-essential amino acids (ml)* |  | 15.128 |
| Sodium Glutamate (ml) | 100 g/L | 3.78 |
| Vitamin (ml)* | 47.6x | 21 |
| Folic Acid (ml) | 0.5 g/L | 1 |
| Propionic Acid (ml) | - | 6 |
| Nipagin (ml)* | 100 g/L | 15 |

\* Refer to Piper et al., 2014 for complete composition

